# Mieap forms membrane-less organelles to compartmentalize and facilitate cardiolipin metabolism

**DOI:** 10.1101/2020.10.26.354365

**Authors:** Naoki Ikari, Katsuko Honjo, Yoko Sagami, Yasuyuki Nakamura, Hirofumi Arakawa

## Abstract

Biomolecular condensates (BCs) are formed by proteins with intrinsically disordered regions (IDRs) via liquid–liquid phase separation. Mieap/Spata18, a p53-inducible protein, participates in suppression of colorectal tumors by promoting mitochondrial quality control. However, the regulatory mechanism involved remains unclear. Here, we report that Mieap is an IDR-containing protein that drives formation of BCs involved in cardiolipin metabolism. Mieap BCs specifically phase separate the mitochondrial phospholipid, cardiolipin. Mieap directly binds to cardiolipin *in vitro*. Lipidomic analysis of cardiolipin suggests that Mieap promotes enzymatic reactions in cardiolipin biosynthesis and remodeling. Accordingly, four cardiolipin biosynthetic enzymes, TAMM41, PGS1, PTPMT1, and CRLS1, and two remodeling enzymes, PLA2G6 and TAZ, are phase-separated by Mieap BCs. Mieap-deficient cells exhibit altered crista structure, leading to decreased respiration activity and ATP production in mitochondria. These results suggest that Mieap may form membrane-less organelles to compartmentalize and facilitate cardiolipin metabolism, thus potentially contributing to mitochondrial quality control.

## Introduction

BCs in cells are also known as liquid droplets because of their liquid-like nature. BCs are composed of proteins, nucleic acids, and other macromolecular components^1–3^, and they are formed by proteins with intrinsically disordered regions (IDRs) via liquid–liquid phase separation (LLPS). Importantly, BCs function as membrane-less organelles (MLOs) that compartmentalize and facilitate cellular biological reactions. BCs are not surrounded by lipid bilayers, which enables facile exchange of reactants and products with their surroundings, expediting biological reactions in cells. Furthermore, theoretically, BCs are able to appear and disappear within a cell in response to cellular stress and/or subcellular circumstances, exhibiting spatiotemporally dynamic properties. On the basis of these features, the concept of MLOs is fundamentally different from the long-standing concept of membrane-bound organelles, such as nuclei, lysosomes, mitochondria, and endoplasmic reticulum, stable structures that segregate biomolecules and biological reactions using lipid bilayer membranes. Multiple MLOs may exist within a cell, possibly regulating cellular biological reactions and activities, including transcriptional regulation^4^, signal transduction^5, 6^, immunity^7^, centrosome activity^8^, and mitosis^9^. This list is rapidly growing.

Metabolic reactions are believed to be highly organized through spatiotemporal clustering and compartmentalization of sequential enzymes and substrates/intermediates at subcellular sites, which maximizes efficiency of linked reactions^10^. Without this mechanism, small, toxic intermediates, derived from metabolic reactions, rapidly diffuse throughout the cytoplasm. Thus, the concept of subcellular compartmentalization of metabolic reactions has been anticipated for a long time. It was initially conceived as a “metabolon,” a structural-functional complex of sequential metabolic enzymes and substrates/intermediates^11^. The “metabolon” concept predicted that sequential enzymes and cellular structural elements form a supramolecular complex for metabolic reactions. In fact, many studies have demonstrated that enzyme clustering *in vivo* may facilitate sequential enzymatic metabolic reactions^12–15^. Among them, “purinosomes” involved in purine metabolism, were the first demonstration of enzyme clustering, revealed by live-cell imaging of six sequential enzymes for *de novo* purine synthesis^16^. Although MLOs that compartmentalize and expedite metabolic reactions in cells could address regulatory mechanisms for sequential metabolic reactions, there has been no clear evidence for metabolic MLOs until now.

Cardiolipin (CL) is an important phospholipid for the following reasons. [1] It is a phospholipid dimer, with two phosphate residues and four fatty acyl chains^17^. [2] Because of this unique structure, it forms a cone shape, contributing to curvature of the lipid membrane and to maintenance of mitochondrial cristae^18^. [3] CL is the only phospholipid that is specific to mitochondria and is mainly located at the inner mitochondrial membrane and contact sites^18^. [4] CL interacts with many mitochondrial membrane proteins, including electron transport chain complexes involved in oxidative phosphorylation, and the ADP/ATP carrier. Interaction with CL promotes activity of these proteins^18^. [5] CL stabilizes the structural assembly and activity of respiratory super-complexes at the inner mitochondrial membrane^19^. [6] It interacts with cytochrome *c* at the outer surface of the inner mitochondrial membrane, which supports cellular viability by maintaining stable respiratory ATP production and inducing apoptosis, respectively^19^. Therefore, control of CL quality and quantity appears to be important for various mitochondrial functions.

Altered CL metabolism and/or deterioration of CL quality and quantity due to CL metabolic enzyme deficiency causes various mitochondrial dysfunctions in eukaryotes ranging from yeast to mammals^20, 21^. Specifically, Barth syndrome is a human disease caused by mutations in the TAZ gene encoding tafazzin, the enzyme responsible for CL remodeling^22^. Clinical symptoms of Barth syndrome patients include cardiomyopathy, skeletal muscle weakness, neutropenia, and growth retardation. Barth syndrome models, including lymphoblasts and induced pluripotent stem cells derived from patients, exhibit abnormal crista structures, decreased respiration activity, and increased reactive oxygen species (ROS) generation in mitochondria^23, 24^. This implies that substantial regulation of CL metabolic reactions is required to maintain mitochondrial functions and to prevent diseases. However, the regulatory mechanism for CL metabolic reactions is unknown.

Mitochondria-eating protein (Mieap, also denominated SPATA18) was originally identified as a p53-inducible protein. Its mRNA expression is directly regulated by the tumor suppressor, p53, in response to various cellular stresses, including DNA damage^25^. Mieap expression is lost in nearly 50% of human cancer cell lines due to promoter methylation. Mieap-deficient LS174T colorectal cancer cells generate higher levels of mitochondrial ROS and synthesize less ATP. Mitochondrial ROS in Mieap-deficient colorectal cancer and gastric cancer cells enhances migration and invasiveness of cancer cells under hypoxic conditions^26, 27^. Mieap deficiency promotes intestinal tumors in Apc^Min/+^ mice^28^. Tumors in Mieap-deficient ApcMin/+ mice reveal abnormal mitochondrial morphology, such as large size, round shape, and disordered cristae. Mieap expression is defective in thyroid oncocytic cell tumors, which accumulate abnormal mitochondria in tumor cells^29^. Mieap-regulated mitochondrial quality control is inactivated in tumor tissues of nearly 70% of colorectal cancer and 25% of breast cancer patients^26, 30^. These observations suggest that Mieap suppresses tumors via mitochondrial quality control. However, the mechanism underlying regulation of mitochondrial quality control by Mieap remains unclear.

Here, we report that Mieap is an IDR-containing protein that drives BC formation in mitochondria. Mieap BCs phase separate the mitochondrial phospholipid, CL. The present findings suggest that Mieap BCs function as MLOs that compartmentalize and promote CL synthesis and remodeling reactions, leading to stabilization of oxidative phosphorylation and suppression of mitochondrial ROS generation. Thus, we suggest that the Mieap-CL axis is the regulatory mechanism for efficient CL metabolic reactions and/or CL quality and quantity control, forming MLOs that govern mitochondrial quality control by regulating CL synthesis and remodeling. Dysregulation of this pathway leads to mitochondrial dysfunction due to CL alterations, causing a variety of diseases and/or pathologies, including cancer and obesity.

## Results

### Mieap forms biomolecular condensates in mitochondria

Mieap has been reported to form vacuole-like structures, designated as Mieap-induced vacuoles (MIVs)^31^. To confirm this hypothesis, we performed the IF experiment on structures comprising EGFP-Mieap. EGFP-Mieap reproducibly formed green condensates, while antibodies (anti-Mieap antibody and anti-GFP antibody) produced ring-shaped staining around the green condensates (Figure S1A). Thus, we suspected that antibodies are unable to permeabilize EGFP-Mieap condensates.

We performed additional IF using anti-FLAG antibody for the MIV structures, which comprise N-FLAG-Mieap (N-terminal FLAG-tagged Mieap protein) and C-FLAG-Mieap (C-terminal FLAG-tagged Mieap protein). Ring-shaped staining was observed with N-FLAG-Mieap, but not C-FLAG-Mieap (Figure S1B), suggesting that Mieap may be positioned with its N-terminal domain facing outward at the surface of the condensates.

Using transmission electron microscopy (TEM), MIV structures appeared to consist of two phases: a dominant electron-dense phase that stained strongly with OsO_4_, and an OsO_4_-negative, electron-lucent minor phase (Figure S1C). In immunoelectron microscopy (IEM) analysis with anti-Mieap antibody, gold colloid staining indicated that Mieap protein was distributed over the major electron-dense phase (Figure S1D). Thus, both fluorescent tagging and antigen-antibody reactions confirmed that Mieap forms protein condensates.

Mieap condensates exhibited spherical or oval shapes, fusion, and multi-phase structure consisting of two phases: a Mieap-containing phase and a Mieap-depleted phase (Figure S1A, S1C, S1D and Movie S1). These characteristics, and their propensity to fuse, are not contradictory to a notion that Mieap condensates have liquid-like properties, suggesting that these structures are droplets^3^. Therefore, we designate Mieap-induced structures as Mieap BCs (Mi-BCs).

To clarify the spatial relationship between Mi-BCs and mitochondria, we performed live-cell imaging using cells co-expressing EGFP-Mieap and an outer mitochondrial membrane marker, mApple-TOMM20. mApple-TOMM20 entirely surrounded Mi-BCs, suggesting that Mieap forms condensates in mitochondria (Figure 1, and Movie S2).

**Figure 1.**
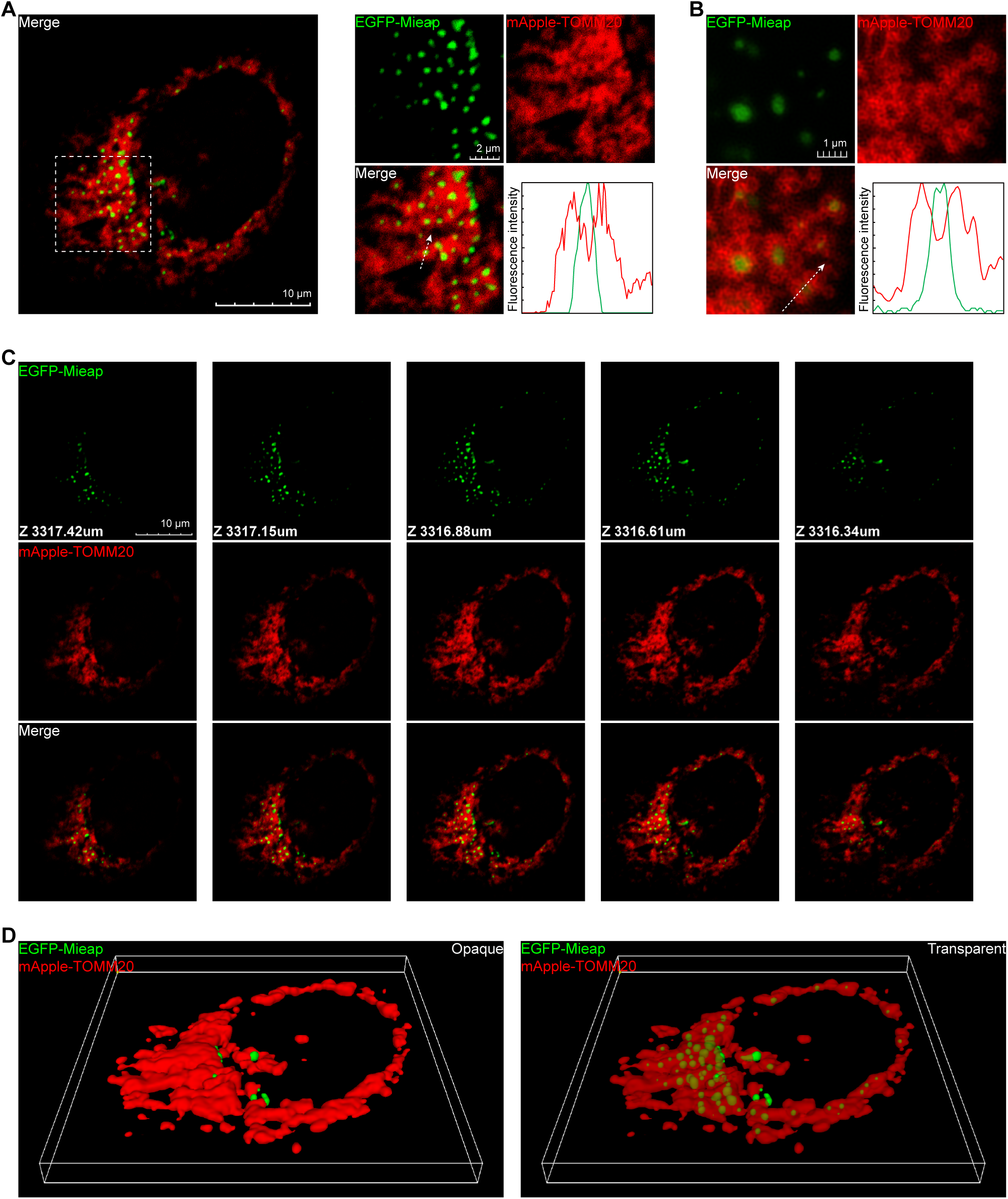
Mieap forms biomolecular condensates in mitochondria. (**A**) Confocal microscopy showing the spatial relationship between Mi-BCs (EGFP-Mieap) and mitochondrial outer membranes visualized with mApple-TOMM20. Left: a cell image. Right: higher magnification of the area indicated by the dashed square in the left panel and a line-scan of fluorescence intensities along the dashed arrow. See also Movie S2. (**B**) Super-resolution images showing the spatial relationship between Mi-BCs (EGFP-Mieap) and mitochondrial outer membranes, visualized with mApple-TOMM20. (**C**) Z-stack images of the cell in (**A**). (**D**) 3D reconstruction of the cell shown in (**A**). See also Movie S2.

### Mieap is an IDR-containing protein with potential to drive LLPS

We performed *in silico* sequence analyses. Mieap orthologs were found in eukaryotes, but not in bacteria, archaea, or viruses (Figure 2A)^32^. Hence, the Mieap function evolved in eukaryotes. Moreover, among eukaryotes, Mieap orthologs were found in metazoans, but not in fungi (Figure 2A), suggesting that Mieap is beneficial to multicellular organisms.

**Figure 2.**
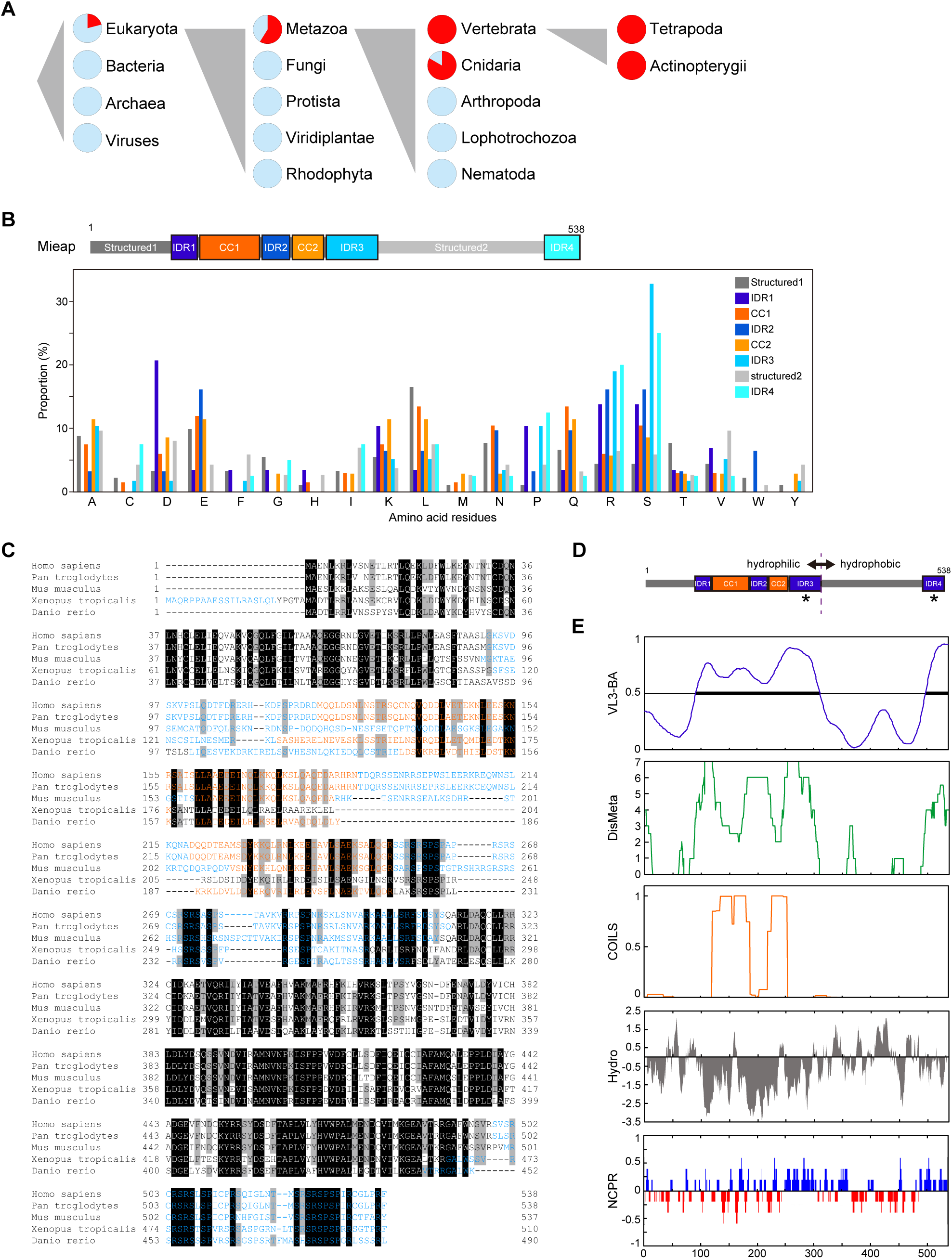
Mieap is an IDR-containing protein. (**A**) Phylogenetic spread of Mieap orthologs annotated with OrthoDB v10. Red sectors indicate present species. Light blue sectors indicate missing species. (**B**) Proportion of amino acid residues in each domain of the Mieap protein. (**C**) Multiple sequence alignment for Mieap orthologs in representative eukaryotes. Black and gray boxes indicate 100% and 80% identical residues among eukaryotes, respectively. Blue letters indicate IDRs annotated by VL3-BA. Orange letters indicate coiled-coil regions annotated by COILS. (**D**) Schematic of the domain structure of Mieap. The dashed vertical line indicates the boundary of gross hydrophilic and hydrophobic halves, separated by IDR3 and the adjacent structured region. Asterisks indicate clusters of positively charged residues. (**E**) Sequence analyses of Mieap protein. VL3-BA prediction of IDRs on the amino acid sequence of Mieap, in which bold lines indicate IDRs; DisMeta, meta-prediction of IDRs on the amino acid sequence of Mieap; COILS; coiled-coil regions annotated on the amino acid sequence of Mieap using a 21-residue sliding window; Hydro, hydrophobicity of Mieap using a 9-residue sliding window; NCPR, the linear net charge per residue of Mieap using a 5-residue sliding window.

Using prediction tools^33–35^, we determined that Mieap has four IDRs occurring around two coiled-coil (CC) domains (amino acids 92–311: IDR1 to IDR3) and in the C-terminus (amino acids 499–538: IDR4) (Figure 2B–E), suggesting that Mieap can potentially drive LLPS^3^.

All four IDRs are enriched in arginine (R, positively charged) and serine (S, uncharged polar) (Figure 2B). Additionally, IDR1 is enriched in aspartic acid (D, negatively charged), and IDR2 is enriched in glutamic acid (E, negatively charged). D and E were mixed with positively charged residues in IDR1 and IDR2, respectively. In contrast, IDR3 and IDR4 formed clusters of positively charged residues, characterized by repeats of R and S (Figure 2B–E) ^36^.

Although amino acid sequences of the IDRs are evolutionarily divergent compared to the structured regions, the distribution of IDRs and clusters of positively charged residues in IDR3 and IDR4 are evolutionarily conserved (Figures 2C, 2E, and S2)^37^. In addition, there is an evolutionarily conserved hydropathic character in the Mieap sequence as a whole, the N-terminal half being hydrophilic and the C-terminal half being hydrophobic (Figures 2D, 2E, and S2) ^37^. This implies that Mieap protein may act as a cellular biosurfactant.

These molecular features in Mieap IDRs are consistent with the concept of the “evolutionary signature” that Zarin et al. previously proposed^38^. They found that although the amino acid sequences of IDRs are poorly conserved in alignment, most disordered regions contain multiple molecular features that are preserved as an “evolutionary signature”, which can be used to predict IDRs from their amino acid sequences in yeast.

### Material state and dynamics of Mi-BCs are determined by specific regions of Mieap

To map sites responsible for the physical state and dynamics of Mi-BCs, using a confocal microscope, we examined cells expressing EGFP-Mieap WT (WT) and three deletion-mutant forms, EGFP-Mieap ΔCC, Δ275, and Δ496 (ΔCC, Δ275, and Δ496) (Figures 3A).

**Figure 3.**
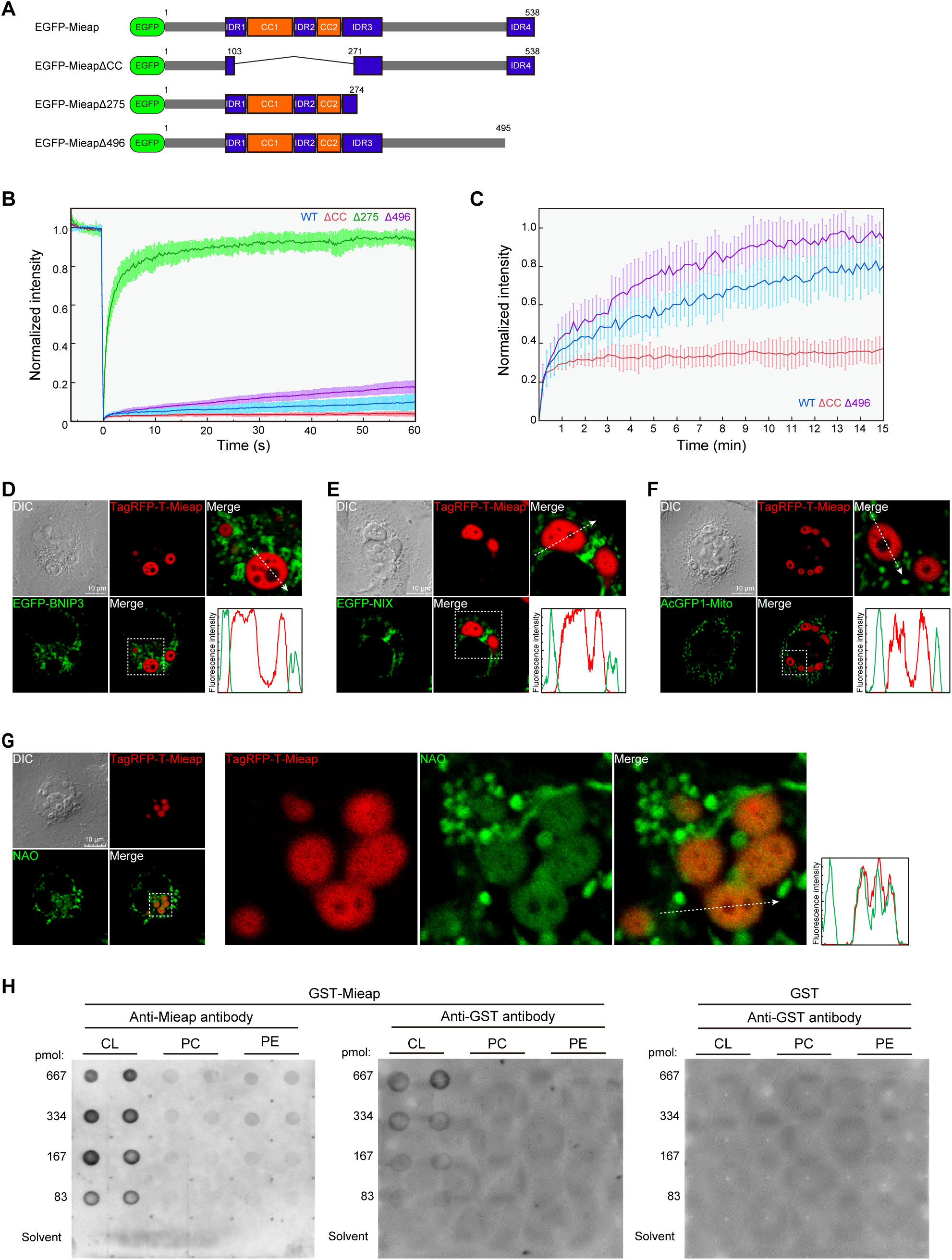
Mi-BCs phase-separate NAO, and Mieap binds to CL. (**A**) EGFP-Mieap and the three deletion-mutant forms. The schematic indicates wild-type (WT) and three deletion mutants (ΔCC, Δ275, and Δ496) of EGFP-Mieap protein. Numbers indicate amino acid residues. (**B**) Normalized average fluorescence recovery in the FRAP experiment. Condensates formed by WT, ΔCC, Δ275, and Δ496 were analyzed in A549 cells. Each condensate was subjected to spot-bleaching using a 488-nm laser at 10% laser power with 11.6 µs/µm exposure time and followed up for 60 s. n = 15 condensates for each construct. Data shown are means ±SD. (**C**) Normalized average fluorescence recovery in the FRAP experiment with weaker laser exposure as in (**B**). Laser power was weakened to 1.4% and exposure time was shortened to 1.4 µs/µm. Observation duration was expanded to 15 min after photobleaching entire condensates. n = 10 condensates for each construct. Data shown are means ±SD. (**D-F**) Live-cell imaging of Mi-BCs and various mitochondrial fluorescence probes in A549 cells. Whether each mitochondrial fluorescence probe is phase-separated by Mi-BCs was examined with live-cell imaging analysis in A549 cells. EGFP-BNIP3 (**a**), EGFP-NIX (**b**), AcGFP-mito (**c**) were not incorporated into Mi-BCs. Lower right: line-scan of fluorescence intensities along the dashed arrow. See also Movie S3. (**G**) Live-cell imaging of Mi-BCs and 10-nonylacridine orange bromide (NAO) in A549 cells. NAO was incorporated into Mi-BCs. Lower right: line-scan of fluorescence intensities along the dashed arrow. See also Movie S4. (**H**) Lipid-binding analysis of GST-tagged Mieap protein. GST-Mieap or GST was incubated with membranes on which increasing amounts of CL, phosphatidylcholine (PC), and phosphatidylethanolamine (PE), ranging from 0-667 pmol, were spotted. Protein-lipid interactions were visualized using an anti-Mieap antibody and/or anti-GST antibody, as indicated.

To investigate protein dynamics in Mi-BCs, we performed fluorescence recovery after photobleaching (FRAP) studies for WT, ΔCC, Δ275, and Δ496 condensates. During observations up to 60 s after spot-bleaching, with the bleaching depth being 82.2±8.1%, fluorescence intensity of WT, ΔCC, Δ275, and Δ496 condensates recovered to 9.9±4.0%, 4.0±1.0%, 94.3±4.0%, and 17.8±3.1% of their initial values, respectively (Figures 3B and S3A). Fluorescence recovery of the Δ275 condensates almost achieved their initial value within the 60-s observation period, indicating the most fluid state.

When we performed less intense laser exposure, which reduced the bleaching depth to 21.2±4.4%, fluorescence recovery increased 60 s after spot-bleaching (WT, 50.4±8.8%; ΔCC, 32.6±8.8%; Δ275, 94.9±13.7%; Δ496, 61.5±10.1%) (Figure S3B). The fluorescence recovery rate increased when the number of bleached molecules was small, suggesting that availability may be a rate-limiting factor.

We further examined slow fluorescence recovery up to 15 min. WT and Δ496 condensates showed continuous fluorescence recovery, which reached 80.4±10.2% and 94.0±9.5% of the initial value within 15 min, respectively (Figure 3C). In contrast, ΔCC condensates reached equilibrium at 37.4±6.3% of their initial value within 15 min (Figure 3C). These data suggested that WT condensates, as well as Δ496 condensates, consist mainly of mobile materials, but protein availability from their surroundings is limited. Therefore, FRAP analysis data suggested that Mi-BCs exist in mitochondria.

### Mi-BCs phase-separate the mitochondrial phospholipid, CL

To identify molecules targeted for phase separation by Mi-BCs, we screened available fluorescence-tagged mitochondrial proteins and mitochondrial fluorescence probes using confocal live-cell imaging. EGFP-BNIP3, EGFP-NIX, AcGFP1-Mito, DsRed2-Mito, and SYBR Green I (a probe for mitochondrial DNA) were localized at mitochondria, but none of them were incorporated into to Mi-BCs (Figure 3D–F and Movie S3).

However, 10-N-nonyl acridine orange (NAO) were specifically incorporated into Mi-BCs (Figure 3G and Movie S4). NAO targets CL^39^. CL binds to >60 mitochondrial proteins via its hydrophobic and electrostatic interactions^40^. As described above, positively charged residues are enriched in Mieap IDRs and Mieap has its C-terminal half hydrophobic region. CL carries two negative charge phosphate residues and four hydrophobic fatty acyl chains ^41^. Sequence data reveal the amphiphilicity of Mieap, and Mi-BCs consist of an electron-dense phase that is positive for OsO_4_ ^42^, according to TEM analysis, suggesting that Mi-BCs may contain unsaturated lipids. Therefore, we suspected that CL is a bona-fide target for phase separation by Mi-BCs.

To determine whether Mieap binds directly to CL, we performed a fat blot assay^43^, in which binding of GST-Mieap to CL, phosphatidylcholine (PC), and phosphatidylethanolamine (PE) was evaluated on lipid-dotted membranes. GST-Mieap bound to CL, but not to PC or PE (Figure 3H).

### Mi-BCs compartmentalize and concentrate CL synthetic and remodeling enzymes

We performed mass spectrometric analyses of CL in A549 cells with and without enforced expression of exogenous Mieap protein by Ad-Mieap infection. The total amount of CL per cell was higher in Ad-Mieap infected cells than in cells without the infection (Figures 4A). Broad CL species showed higher absolute values in cells infected with Ad-Mieap than in non-infected cells (Figures 4B). In contrast, relative amounts of most CL species did not change. However, Mieap significantly increased the proportions of CL72:5, CL72:6, CL70:6, CL68:5, and CL68:6 (Figures 4B). These results suggest that Mieap is involved in CL metabolism. Therefore, we speculated that Mi-BCs may function as MLOs to compartmentalize and facilitate CL metabolic reactions. To validate this hypothesis, we examined whether Mi-BCs phase-separate enzymes sequentially involved in CL metabolism (Figures 4C)^20, 44, 45^.

**Figure 4.**
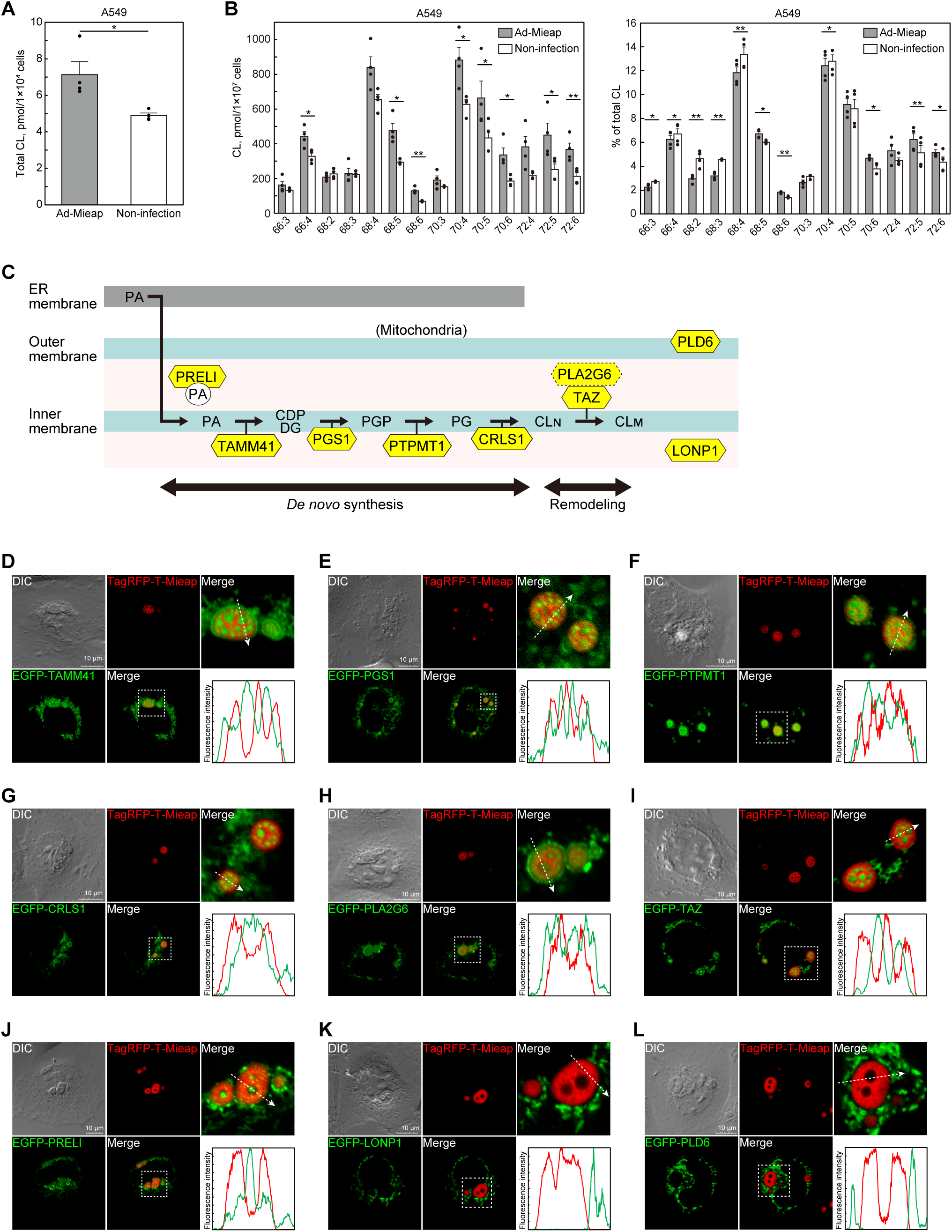
Mi-BCs compartmentalize and concentrate CL synthetic and remodeling enzymes. (**A**) Quantitative assessment of total CL by mass spectrometric analysis. Uninfected A549 cells or cells infected with Ad-Mieap were subjected to mass spectrometric analysis for CL. Data shown are means ±SE (n = 4). Statistical significance is indicated by * (p < 0.05). (**B**) Quantitative and rate assessments of CL species by mass spectrometric analysis. A549 cells were analyzed as described in (**A**). Absolute values of selected CL species are shown as amounts of substance per cell (left panel). Relative values of selected CL species are shown as % of total CL (right panel). Data shown are means ±SE (n = 4). Statistical significance is indicated by * (p < 0.05), or ** (p < 0.01). (**C**) The conventional CL metabolic pathway. Abbreviations: PA, phosphatidic acid; CDP-DG: cytidine diphosphate diacylglycerol; PGP, phosphatidylglycerophosphate; PG, phosphatidylglycerol; CLN, nascent cardiolipin; CLM, mature cardiolipin. (**D – J**) Live-cell imaging of Mi-BCs and CL metabolic enzymes. EGFP-TAMM41 (**D**), EGFP-PGS1 (**E**), EGFP-PTPMT1 (**F**), EGFP-CRLS1 (**G**), EGFP-PLA2G6 (**H**), EGFP-TAZ (**I**), and EGFP-PRELI (**J**) were incorporated into Mi-BCs. Lower right: line-scan of fluorescence intensities along the dashed arrow. See also Movie S5. (**K, L**) Live-cell imaging of Mi-BCs and LONP1/PLD6. EGFP-LONP1 (**K**) and EGFP-PLD6 (**l**) were not incorporated into Mi-BCs. Lower right: line-scan of fluorescence intensities along the dashed arrow. See also Movie S5.

Thus, we examined involvement of the following EGFP-tagged enzymes required for CL metabolism in Mi-BCs by performing confocal live-cell imaging: EGFP-TAMM41, EGFP-PGS1, EGFP-PTPMT1, and EGFP-CRLS1 (involved in CL biosynthesis); EGFP-PLA2G6 (related to CL hydrolysis by phospholipase A_2_ activity); and EGFP-TAZ (involved in CL remodeling)^20, 45^. All of these enzymes localized at mitochondria and were subsequently incorporated into Mi-BCs (Figures 4D–4I and S4A–S4F, and Movie S5). Interestingly, all of these CL metabolic enzymes tended to be concentrated in the Mieap-depleted phase in Mi-BCs (Figures 4D–4I and Movie S5).

We further examined three additional mitochondrial proteins, PRELI (EGFP-PRELI), mitochondrial protease LONP1 (EGFP-LONP1), and mitochondrial CL hydrolase/mitochondrial phospholipase (MitoPLD)/phospholipase D6 (EGFP-PLD6) (Figure S4G-S4I). PRELI is a mitochondrial carrier of PA for CL production^44^. LONP1 is a AAA mitochondrial protease^46^. PLD6 hydrolyzes CL to generate PA at the outer mitochondrial membrane^47^. EGFP-PRELI was incorporated into Mi-BCs (Figures 4J and Movie S5), suggesting that Mi-BCs can also be supplied with PA as a substrate for CL synthesis. Both EGFP-LONP1 and EGFP-PLD6 were located at mitochondria, but neither was incorporated into Mi-BCs (Figure 4K and 4I, and Movie S5).

### Mieap generates the multi-phase structure of Mi-BCs via both N-terminal hydrophilic and C-terminal hydrophobic regions

Using EGFP-TAMM41 and NAO, we determined whether ΔCC, Δ275 and Δ496 BCs are located inside mitochondria by performing live-cell imaging analysis with tomographic 3D reconstruction.

Mi-BCs are fully enveloped by EGFP-TAMM41, signals of which are continuously localized from tubular mitochondria to all around the surfaces of spherical Mi-BCs (Figure 5A and Movies S6 and S8). We further confirmed that NAO is also continuously localized from tubular mitochondria, all round and inside of spherical Mi-BCs (Figure 5E, and Movies S7 and S8).

**Figure 5.**
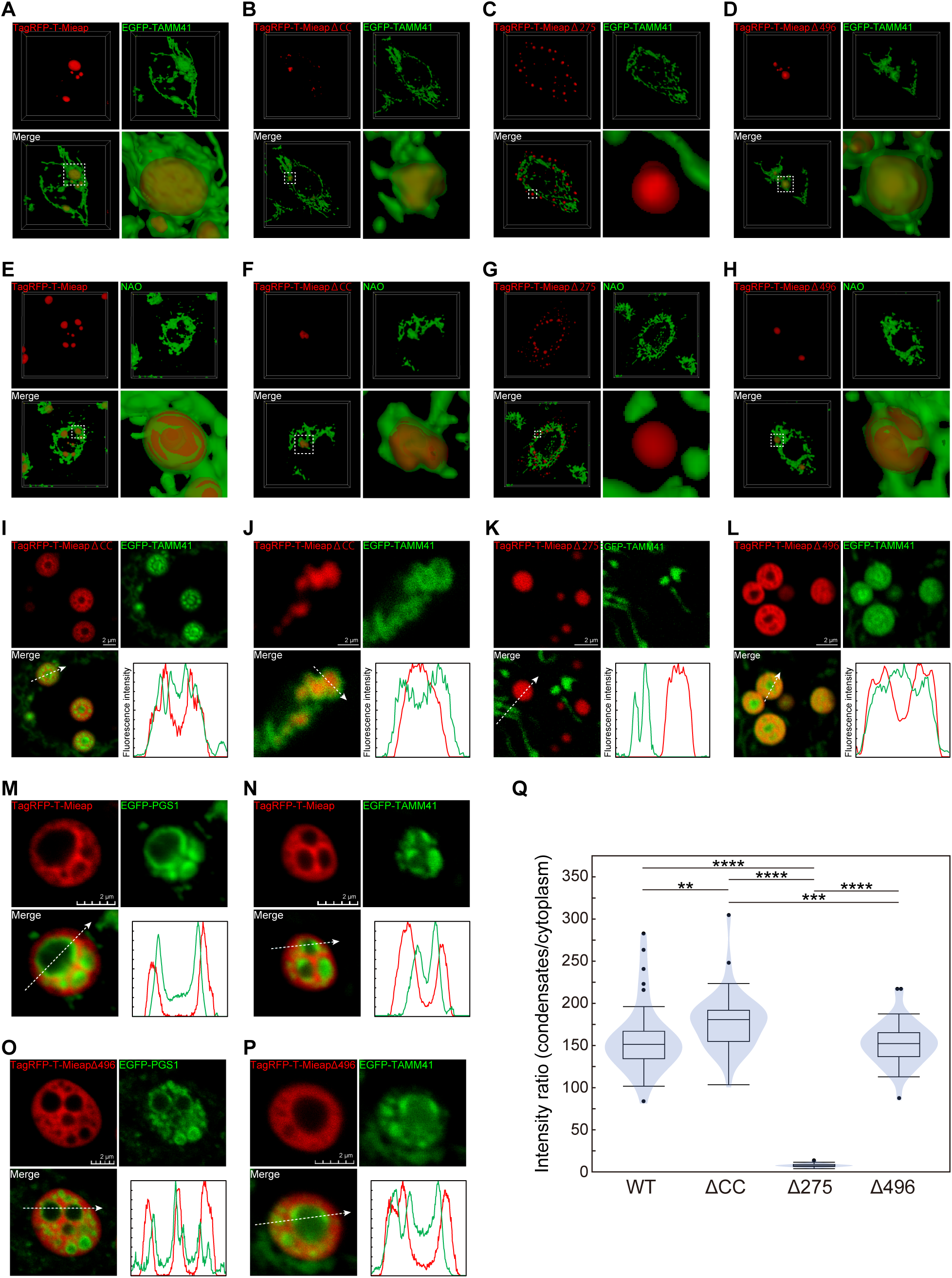
Both N- and C-terminal regions of Mieap are required to generate the multi-phase structure of Mi-BCs. (**A – D**) 3D reconstruction showing the spatial relationship between BCs formed by TagRFP-T-Mieap (**A**), ΔCC (**B**), Δ275 (**C**), and Δ496 (**D**) and mitochondrial inner membranes visualized with EGFP-TAMM41 in HeLa cells. See also Supplementary Movies 6 and 8. (**E – H**) 3D reconstruction showing the spatial relationship between BCs formed by TagRFP-T-Mieap (**E**), ΔCC (**F**), Δ275 (**G**), and Δ496 (**H**) and mitochondrial inner membranes visualized with NAO in HeLa cells. See also Supplementary Movies 7 and 8. (**I** – **L**) Comparison of phase-separating behaviors on the CL metabolic enzyme, TAMM41 (EGFP-TAMM41), between BCs formed by TagRFP-T-Mieap WT (**I**), ΔCC (**J**), Δ275 (**K**), and Δ496 (**L**) in HeLa cells. Right: line-scan of fluorescence intensities along the dashed arrow. (**M, N**) CL metabolic enzymes wet the interface in the Mieap-depleted phase of Mi-BCs. Distributions of the CL metabolic enzymes, EGFP-PGS1 (**M**) and EGFP-TAMM41 (**N**), in the Mieap-depleted phase of Mi-BCs are shown in HeLa cells. Lower right: line-scan of fluorescence intensities along the dashed arrow. (**O, P**) CL metabolic enzymes wet the interface in the Mieap Δ496-depleted phase of Δ496-BCs. Distributions of the CL metabolic enzymes, EGFP-PGS1 (**O**) and EGFP-TAMM41 (**P**), in the Mieap Δ496-depleted phase of BCs formed by the Δ496 mutant are shown in HeLa cells. Lower right: line-scan as in (M, N). (**Q**) The intensity ratio (IR) of EGFP-Mieap WT, ΔCC, Δ275, or Δ496 protein in condensates and cytoplasm, displayed in violin plot. n = 40 cells for each construct in A549 cells. Statistical significance is indicated by ** (p < 0.01), *** (p < 0.001), or **** (p < 0.0001). When the data were visualized using violin plots, box plots were overlaid. The center line in the box indicates the median. The bottom and top of the box indicate the 25^th^ and 75^th^ percentiles. The whiskers extend 1.5 times the interquartile range (IQR) from the top and bottom of the box unless the minimum and maximum values are within the IQR. The values which fall above or below the whiskers are plotted individually as outliers.

Similarly, both ΔCC and Δ496 BCs are also fully enveloped by EGFP-TAMM41, whose signals are continuously derived from tubular mitochondria (Figure 5B and 5D, and Movies S6 and S8). Moreover, ΔCC and Δ496 BCs are also stained by NAO as a clear picture of tubular mitochondria and each condensate (Figure 5F and 5H, and Movies S7 and S8). However, Δ275 condensates are never related to the signals of EGFP-TAMM41 or NAO (Figure 5C and 5G, and Movies S6–S8). These results suggest that ΔCC and Δ496 BCs are located inside mitochondria, whereas Δ275 BCs are present outside mitochondria.

To explore the mechanism responsible for multi-phase structure in Mi-BCs, using EGFP-TAMM41, we examined whether CL metabolic enzymes are phase-separated by ΔCC, Δ275, and Δ496 condensates (Figure 5I – 5L). As expected, Δ275 condensates did not phase-separate EGFP-TAMM41 (Figure 5K). Importantly, although both ΔCC and Δ496 are located in mitochondria, EGFP-TAMM41 was phase-separated and incorporated in the Mieap-depleted phase of Δ496 condensates (Figure 5I), whereas EGFP-TAMM41 was mainly localized across the surfaces of ΔCC condensates, which did not generate multi-phase structures (Figure 5J). These results suggest that both the N-terminal hydrophilic and C-terminal hydrophobic regions are required to form multi-phase droplets. The region of IDR1-3 and two CCs may be critical to the interaction with CL metabolic enzymes to generate the internal enzyme-containing phase (the Mieap-deficient phase) in Mi-BCs.

We further explored the internal structure of the Mieap-depleted phase with EFGP-PGS1 or EGFP-TAMM41. Importantly, we found that CL biosynthetic enzymes, EGFP-PGS1 or EGFP-TAMM41, formed condensates in the Mieap-depleted phase, which wetted the interface between the Mieap-containing phase and the Mieap-depleted phase in either WT or Δ496 BCs (Figure 5M – 5P). These results suggest that enzymatic reactions between CL metabolic enzymes and their substrates may occur at the interface between the Mieap-containing phase (Mieap and substrates) and the Mieap-depleted phase (CL metabolic enzymes).

Finally, we examined partitioning behaviors of WT, ΔCC, Δ275, and Δ496 proteins by performing analysis of Intensity Ratios of each protein in BCs and cytoplasm^48^. Intensity Ratios values (condensates/cytoplasm) of WT, ΔCC, and Δ496 are 158.62±40.74, 178.81±34.07, 8.29±1.92, and 153.14±25.66 (mean±SD) (Figure 5Q), respectively. This implies that Intensity Ratios of WT, ΔCC, and Δ496 are more than 18 times higher than that of Δ275. All WT, ΔCC, and Δ496 BCs are localized within mitochondria, whereas Δ275 condensates are present in cytoplasm. Therefore, these results suggest that Mieap protein tends to be highly partitioned and concentrated in mitochondrial BCs, and that this propensity is possibly determined by the C-terminal hydrophobic region of Mieap, which could mediate interaction of Mieap with CL/CL-related phospholipids.

### Mieap functions in mitochondrial quality control via regulation of CL metabolism

CL alterations cause mitochondrial dysfunction^20, 21^. Mieap is thought to be involved in mitochondrial quality control^25, 26, 28, 31^. Therefore, we hypothesized that Mieap contributes to mitochondrial quality control by regulating CL metabolism. To test this hypothesis, we evaluated mitochondrial status relative to CL integrity in the presence or absence of Mieap protein in cells and mice.

First, we examined respiration rate, mitochondrial ATP production rate, crista morphology, and ROS levels, in control and Mieap-knockdown (KD) LS174T cells under physiological conditions, all of which reflect CL integrity. Flux analysis indicated that respiration and ATP production rates of Mieap-KD cells were significantly lower than those of control cells (Figure 6A and 6B). TEM analysis revealed that cristae of Mieap-KD cells decreased, and their morphology became indistinct and irregular, compared to that of control cells (Figure 6C and 6D). ROS levels increased in Mieap-KD cells (Figure S5). Consistently, the total amount of CL in control cells was higher than in Mieap-KD cells (Figure 6E), and control cells showed higher absolute values for CL species than Mieap-KD cells (Figure 6F). Furthermore, physiological Mieap significantly increased the relative values of CL72:4, CL72:5, CL70:4, CL68:3, and CL68:4 in LS174T cells (Figure 6F).

**Figure 6.**
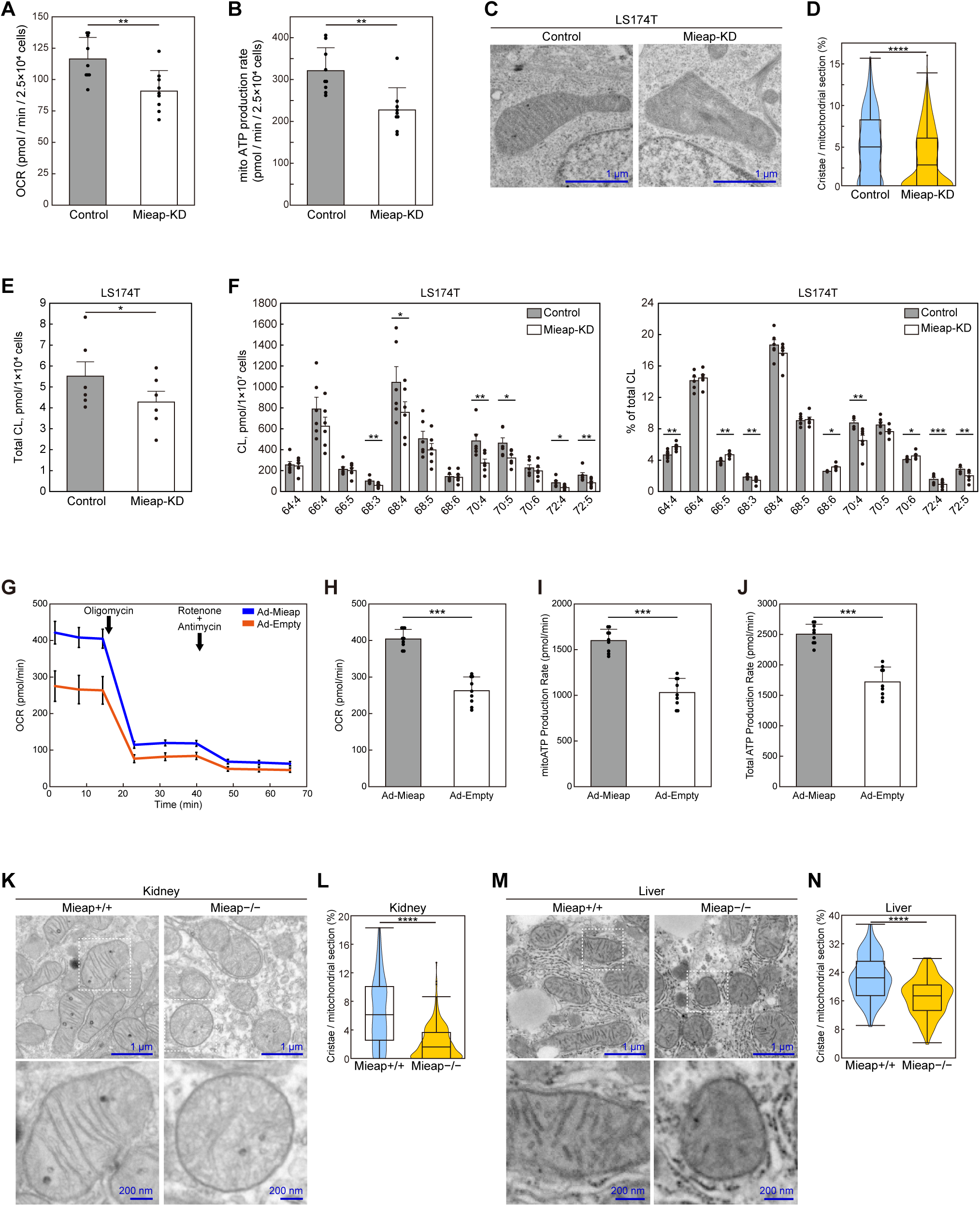
Mieap contributes to mitochondrial quality control by promoting CL metabolism. (**A**) Oxygen consumption rates (OCR) of LS174T-cont and Mieap-KD cells under normal conditions calculated with a flux analyzer. Data are shown as means ± SD (n = 9). (**B**) Mitochondrial ATP production rates of LS174T-cont and Mieap-KD cells under normal conditions calculated with a flux analyzer, using a Seahorse XF real-time ATP rate assay. Data are shown as means ± SD (n = 9). (**C**) Morphology of mitochondria of LS174T-cont and Mieap-KD cells with transmission electron microscopy (TEM). (**D**) Ratio of crista area per mitochondrial section of LS174T-cont and Mieap-KD cells. Quantitative data were obtained from cont mitochondria (n=197) and Mieap-KD mitochondria (n= 329) in TEM images and displayed in a violin plot. (**E**) Quantitative assessment of total CL by mass spectrometric analysis. LS174T cells with (Cont) and without (Mieap-KD) endogenous Mieap expression were subjected to mass spectrometric analysis. Data shown are means ± SE (n = 6). (**F**) Quantitative and rate assessments of CL species by mass spectrometric analysis. LS174T cells were analyzed as described in (**e**). Absolute values of selected CL species are shown as the amount of substance per cell (left panel). Relative values of selected CL species are shown as % of total CL (right panel). Data shown are means ± SE (n = 6). See also Supplementary Data 1. (**G**) The kinetic profile of the OCR using the Seahorse XF Real-Time ATP rate assay in HCT116 cells infected with Ad-Mieap or Ad-empty. (**H – J** Quantitative assessment of OCR (**H**), mitochondrial ATP production rates (**I**), and total ATP production rates (**J**) of the HCT116 cells as in (**G**). Data are shown as means ± SD (n = 9). (**K**) Morphology of kidney mitochondria of Mieap^+/+^ and Mieap^−/−^ mice with TEM. (**L**) Ratios of crista area per mitochondrial section of Mieap^+/+^ and Mieap^−/−^ mouse kidneys. Quantitative data were obtained from Mieap^+/+^ kidney mitochondria (n=190) and Mieap^−/−^ kidney mitochondria (n= 234) in TEM images and displayed in a violin plot. (**M**) Morphology of liver mitochondria of Mieap^+/+^ and Mieap^−/−^ mice with TEM. (**N**) Ratios of crista area per mitochondrial section of Mieap^+/+^ and Mieap^−/−^ mouse livers. Quantitative data were obtained from Mieap^+/+^ liver mitochondria (n=146) and Mieap^−/−^ liver mitochondria (n= 134) in TEM images and displayed in a violin plot. (**A, B, D, E, F, H-J, L, N**) Statistical significance is indicated by * (p < 0.05), ** (p < 0.01), *** (p < 0.001) or **** (p < 0.0001). When the data were visualized using violin plots, box plots were overlaid. The center line in the box indicates the median. The bottom and top of the box indicate the 25th and 75th percentiles. The whiskers extend 1.5 times the interquartile range (IQR) from the top and bottom of the box unless the minimum and maximum values are within the IQR. The values which fall above or below the whiskers are plotted individually as outliers.

Second, utilizing a Mieap-deficient colorectal cancer cell line HCT116, in which the promoter of the *Mieap* gene is completely methylated^25^, we examined whether re-expression of Mieap affects respiration rate and mitochondrial ATP production rate in these cells. As shown in Figure 6G-J, re-expression of Mieap protein by Ad-Mieap infection significantly increased respiration rate and mitochondrial ATP production in HCT116 cells.

Third, we analyzed crista morphology in kidney and liver tissues of Mieap-knockout (KO) mice by performing TEM analysis. In Mieap^−/−^ kidney mitochondria, irregularly dilated lamellar structures without distinct OsO_4_ staining were observed (Figure 6K and 6M). A decrease in normal crista structure was a common characteristic of Mieap^−/−^ mitochondria in the kidney and liver (Figure 6K-6N).

Fourth, we performed a large-scale cross-sectional observation of 1,225 Mieap^+/+^, Mieap^+/−^, and Mieap^−/−^ mice to identify long-term consequences of Mieap deficiency. Average body weights of Mieap^+/−^ and Mieap^−/−^ mice were higher than those of Mieap^+/+^ mice throughout their lives (Figures 7A and S6). Differences were prominent during middle and old age, from 44 to 104 weeks (Figure 7B) (mean value ± SE; Mieap^+/+^ 33.065±0.425 g [n=149], Mieap^+/−^ 34.048±0.302 g [n=295], Mieap^−/−^ 35.090±0.392 g [n=175]), but particularly during middle age, from 53 to 62 weeks (Figure 7C) (mean value ± SE; Mieap^+/+^ 31.048±0.759 g [n=49], Mieap^+/−^ 33.378±0.496 g [n=115], Mieap^−/−^ 34.579±0.645 g [n=68]).

**Figure 7.**
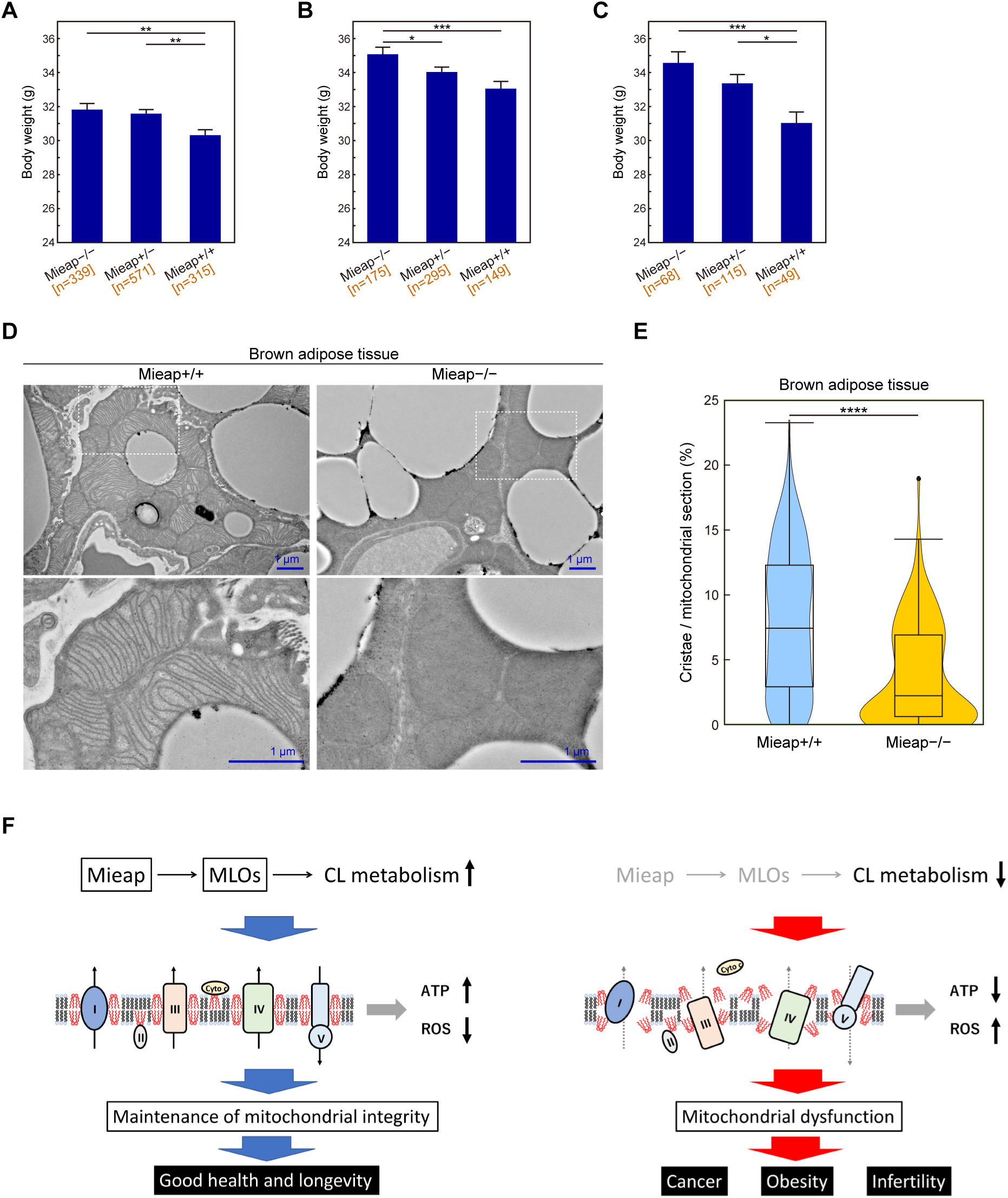
Mieap prevents obesity by maintaining cristae structures of BAT. (**A** – **C**) Body weights of Mieap^+/+^, Mieap^+/−^, and Mieap^−/−^ mice (7-130 weeks of age) (**A**), (44-104 weeks of age) (**B**), and (53-62 weeks of age) (**C**). Data shown are means ±SE. Statistical significance is indicated by * (p < 0.05), ** (p < 0.01), or *** (p < 0.001). (**D**) Morphology of brown adipose tissue (BAT) mitochondria of Mieap^+/+^ and Mieap^−/−^ mice with TEM. (**E**) The ratio of crista area per mitochondrial section of Mieap^+/+;^ and Mieap^−/−^ mice BAT. Quantitative data were obtained from Mieap^+/+^ BAT mitochondria (n=181) and Mieap^−/−^ BAT mitochondria (n=129) in TEM images and displayed in a violin plot. Statistical significance is indicated by **** (p < 0.0001). The center line in the box indicates the median. When the data were visualized using violin plots, box plots were overlaid. The bottom and top of the box indicate the 25^th^ and 75^th^ percentiles. The whiskers extend 1.5 times the interquartile range (IQR) from the top and bottom of the box unless the minimum and maximum values are within the IQR. The values which fall above or below the whiskers are plotted individually as outliers. (**F**) A hypothetical model for mitochondrial quality control via the Mieap-MLOs-CL metabolism axis.

Reduced mitochondrial respiratory activity in adipose tissues has been suggested as a factor contributing to obesity^49^. Brown fat tissue (BAT) is essential to heat production via both its respiratory activity, which generates a proton gradient, and uncoupling protein 1 (UCP1)-mediated proton leakage across the mitochondrial inner membrane^50, 51^. Therefore, BAT mitigates obesity. Importantly, as a CL-binding protein, activity of UCP1 is stabilized by CL^52^. Thus, we examined the status of mitochondrial cristae in BAT of Mieap^−/−^ mice. We confirmed that normal crista structure was significantly decreased in Mieap-deficient BAT (Figure 7D and 7E), suggesting that obesity may be a long-term consequence of mitochondrial dysfunction due to CL alteration in tissues of Mieap-deficient mice, including BAT (Figure 7F).

## Discussion

Although the importance of multi-enzyme complexes in metabolic enzyme reactions has been recognized, it remains unclear how this complex of enzymes efficiently and safely enables sequential enzymatic reactions by preventing diffusion of intermediates. A recent report suggested that concentration of multiple enzymes and substrates/intermediates in a restricted space could mediate efficient sequential enzymatic reactions by preventing diffusion of intermediates^53^. In that model, multiple copies of upstream and downstream enzymes involved in sequential enzymatic reactions are assembled into a single cluster, called an “agglomerate.” According to this model, “once an upstream enzyme produces an intermediate, although the probability of the intermediate being processed by any individual downstream enzymes is low, the probability that the intermediate will be processed by one of the many downstream enzymes in the ‘agglomerate’ can be high.”^53^ Therefore, based on this model, molecular crowding of enzymes, substrates, and intermediates in a restricted space could enable efficient sequential enzymatic reactions mediated by multiple enzymes.

The “agglomerate” concept is promising, but an important question remains. How can so many diverse molecules, including multiple enzymes, substrates, and intermediates, be gathered, concentrated, and compartmentalized in a single restricted space? What drives formation of the “agglomerate”? We speculate that BCs could organize agglomerates as metabolic BCs^54^. Accumulating evidence suggests that BCs function as MLOs, which promote biochemical reactions by concentrating and compartmentalizing enzymes and substrates in cells^55–59^. Since BCs are not surrounded by lipid bilayers, theoretically, they exhibit spatiotemporal dynamic properties within a cell in response to cellular stress and/or subcellular circumstances. More importantly, while BCs contain hundreds of molecules, a few scaffold proteins can drive formation of these MLOs^1^. If there are proteins that can organize metabolic BCs as scaffolds^57^, these agglomerates could enable efficient metabolic reactions.

In the present study, we obtained the following evidence supporting our hypothesis that Mi-BCs may regulate CL metabolism: [1] Mieap drives formation of droplets in mitochondria. [2] Mi-BCs phase-separate NAO, a specific probe for CL, and Mieap directly binds to CL. [3] Mi-BCs compartmentalize and concentrates all four sequential enzymes for CL biosynthesis (TAMM41, PGS1, PTPMT1, and CRLS1). [4] Mi-BCs compartmentalize and concentrates two enzymes for CL remodeling (PLA2G6 and TAZ). [5] The presence or absence of Mieap protein is closely related to an increase or decrease in various species of CL, respectively. [6] Mieap protein specifically increases the proportion of several species of CLs. [7] Mieap deficiency is related to changes in crista structure and CL metabolism in cells, and crista structure *in vivo*. This evidence suggests that Mieap is a scaffold protein that drives formation of metabolic BCs to compartmentalize and concentrate enzymes, substrates, and intermediates that are involved in CL biosynthesis and remodeling, leading to molecular crowding within Mi-BCs that promotes efficient catalysis of CL metabolic reactions.

Mi-BCs exhibit properties of multi-phase droplets, in which there are two phases, a Mieap-containing phase and a Mieap-depleted phase. Interestingly, CL and Mieap occur in the Mieap-containing phase, whereas all CL biosynthesis and remodeling enzymes, including TAMM41, PGS1, PTPMT1, CRLS1, PLA2G6, and TAZ, are predominantly segregated into the Mieap-depleted phase. This result suggests that substrates, intermediates, and products for CL metabolism do not reside in the same phase as their catalytic enzymes. Such a relationship between substrates and enzymes in multi-phase droplets are not limited to Mi-BCs but seen in other droplets. In terms of RNA processing droplets formed by FMRP and CAPRIN1, RNA and phosphorylated FMRP form multi-phase droplets, in which the deadenylation enzyme, CNOT7, and the substrate, polyA-RNA, are segregated into different phases, but this leads to faster deadenylation rates^60^.

Why don’t substrates and enzymes occur in the same phase of metabolic BCs? A recent study demonstrated that sequestration of enzymes to a membrane-less compartment that is away from, but adjacent to substrates, can accelerate reactions much faster than when the enzymes are mixed with the substrates in the same compartment^61^. Concentration of enzymes and substrates in a single phase might result in substrate inhibition^61^. Therefore, separation of enzymes from their substrates via LLPS could facilitate enzymatic reactions by mitigating substrate inhibition. In this case, the interface between the enzyme and substrate phases would be the site of the reaction. Consistent with this hypothesis, we observed accumulation of CL biosynthetic enzymes such as PGS1 and TAMM41 at the interface of the Mieap-depleted phase. Therefore, enzymatic reactions of CL enzymes and CL substrates may occur at the interface between the Mieap-containing phase (Mieap and substrates) and the Mieap-depleted phase (CL metabolic enzymes) (Figure S7).

Feric et al. reported the mechanism for generation of multi-phase structures in droplets^62^. By performing *in vivo* and *in vitro* experiments, the authors demonstrated that layered droplet organization is caused by differences in droplet surface tension, facilitating sequential RNA processing reactions in a variety of RNP bodies. In their experiments, F1B and NPM1 formed multi-phase droplets in which F1B droplets tended to be encapsulated within NPM1 droplets. They found that F1B droplets tended to wet hydrophobic surfaces, whereas NPM1 droplets tended to wet hydrophilic surfaces. Wetting refers to the contact between liquids and surfaces, which depends on surface tension. Therefore, in the aqueous phase, F1B droplets with high surface tension tended to be enveloped by NPM1 droplets with lower surface tension.

Since Mieap may be positioned with its N-terminal domain facing outward at the surfaces of Mi-BCs (Figure S1B), Mi-BCs could exist in a hydrophilic environment. If so, according to the theory of Feric et al., the Mieap-containing phase may be enveloped by the Mieap-depleted phase, because the former is more hydrophobic than the latter. However, the authors also pointed out the presence of a surfactant that modulates surface tension alters or inverts the organization of multi-phase droplets. Since Mieap could serve as a biosurfactant, Mieap may modulate the surface tension of the Mieap-containing phase and the Mieap-depleted phase in a hydrophilic environment. Therefore, the relation between the two phases could be inverted in Mi-BCs.

We hypothesize the following model for sequential CL-metabolic reactions promoted by Mieap (Figure S7). Mieap may stably interact with the substrate (PA) via its C-terminal, hydrophobic, structured region, which exhibits a specific, strong interaction. On the other hand, Mieap weakly and transiently interacts with CL-metabolizing enzymes via its N-terminal hydrophilic region, which exhibits multiple interactions. In this model, once Mieap attracts the substrate with its C-terminal region, Mieap enables sequential CL metabolic reactions by transiently interacting with the enzyme corresponding to the substrate at the interface between the Mieap-containing phase and the Mieap-depleted phase, and then changing enzymes until mature CL is produced. Therefore, interactions of the N-terminal hydrophilic region of Mieap with CL enzymes and the C-terminal hydrophobic region of Mieap with CL/CL-related phospholipids may be critical to drive formation of multi-phase organization of Mi-BCs. In summary, (1) concentration of enzymes and substrates, (2) segregation of enzymes and substrates into distinct sub-compartments of metabolic droplets, (3) interfacial catalysis, and (4) biosurfactant activity of Mieap could foster highly efficient sequential enzymatic reactions for CL metabolism in Mi-BCs. We suggest that Mi-BCs may be the first known metabolic MLOs to regulate CL synthesis and remodeling.

BCs are often thought to accelerate enzymatic reaction rate by merely increasing local concentrations of enzymes and substrates (mass action). However, Peeples and Rosen reported that concentrating enzymes and substrates alone results in decreasing enzymatic reaction rates by substrate inhibition due to high concentrations of substrates^63^. They demonstrated that in addition to mass action, a scaffold-induced decrease in *K*_M_ is critical to accelerate enzymatic reactions in BCs^63^. In their synthetic system where the SUMOylation enzyme cascade is recruited into engineered condensates, they showed that having both enzyme and substrate bound simultaneously to proximal sites in a scaffold oligomer, which leads to decreased *K*_M,_ is required to enhance the enzymatic reaction. Our Mi-BC model is compatible with their findings; however, further investigation of this hypothesis is required.

In the present study, we demonstrated that Mieap-deficient LS174T cells exhibited altered CL metabolism, decreased respiration activity, increased ROS levels, and manifested abnormal crista structure, all of which are consistent with phenotypes induced by CL alteration. We also found that Mieap-deficient mice exhibit decreased numbers and morphological abnormalities of mitochondrial cristae in kidney and liver tissues. Furthermore, our body weight analysis of Mieap^+/+^, Mieap^+/−^, and Mieap^−/−^ mice clarified increased obesity in Mieap-deficient mice, which is likely attributable to mitochondrial dysfunction in various tissues, including BAT. Therefore, we assume that all these phenotypes in Mieap-deficient cells and mice reflect mitochondrial dysfunctions related to abnormal CL metabolism. So far, autophagy and proteostasis are two major mechanisms in mitochondrial quality control^64^. In addition to these, we suggest that the Mieap-regulated pathway is the third mechanism for mitochondrial quality control, in which Mieap maintains integrity of mitochondria by regulating CL metabolism. It accomplishes this through homeostasis of the inner mitochondrial membrane by regulating CL metabolism, stabilizing oxidative phosphorylation, and suppressing mitochondrial ROS generation (Figure 7F).

Previously, we reported that although Mieap-deficient mice did not suffer from intestinal dysfunction, Mieap-deficient Apc^Min/+^ mice exhibited remarkable intestinal tumor generation and malignant transformation, compared to Mieap-WT Apc^Min/+^ mice^28^. Furthermore, mitochondria in Mieap-deficient tumors revealed abnormal morphology, including fewer cristae and enlarged, spherical mitochondria. These results support the role of Mieap in mitochondrial quality control through control of CL metabolism in response to oncogenic stress. In addition to Apc^Min/+^ mice, we also observed tumor suppressive role of Mieap in a thyroid cancer mouse model^65^. Therefore, Mieap-regulated mitochondrial quality control could be critical in tumor suppression by promoting CL metabolism, which leads to upregulation of respiratory activity and downregulation of mitochondrial ROS generation. Considering the role of Mieap in p53 function, we suggest that Mieap could act as a spatiotemporal and dynamic regulator/modulator of CL metabolism to suppress tumor initiation and progression. Recently, we found that Mieap-deficient sperm in Mieap-KO mice cause *in vitro* infertility due to mitochondrial ROS elevation and impaired sperm motility (unpublished data). Therefore, in addition to cancer and obesity, it is possible that alterations of Mieap-regulated mitochondrial quality control also promote infertility (Figure 7F). Further investigation is required to clarify the precise role of Mieap in the regulation of CL metabolism.

## Acknowledgments

We thank H. Suzuki for his contribution to *in vitro* experiments. This work was supported in part by grants from The National Cancer Center Research and Development Fund (29-E-1, 30-A-3) (to HA), JSPS KAKENHI Grant Numbers JP19K07655, JP17H06267, JP20K20305, JP22H02908, and JP22K19412 (to HA), and The Naito Foundation (to HA).

## Author contribution

N.I. and H.A. conceptualization; N.I., K.H., Y.S., Y.N. and H.A. data curation; H.A. funding acquisition; N.I., K.H., Y.S., Y.N. and H.A. formal analysis; N.I. and H.A. investigation; H.A. project administration; H.A. supervision; N.I. and H.A. writing – original draft.

## Materials and Methods

### Cell lines

The following cell lines were purchased from the American Type Culture Collection: A549 (tissue, lung cancer; gender, male), U373MG (tissue, glioblastoma; gender, male), LS174T (tissue, colon cancer; gender, female), and 293 (tissue, embryonic kidney). A549 cells were cultured in RPMI 1640 (Sigma). U373MG, LS174T, and 293 cells were cultured in DMEM (Sigma). All media were supplemented with 10% fetal bovine serum. Cells were maintained at 37°C in a humidified chamber with 5% CO_2_. These cell lines have not been authenticated.

We established a Mieap-KD cell line using LS174T, as previously described^29^. Mieap expression was inhibited in this cell line by retroviral expression of short-hairpin RNA (shRNA) against the Mieap sequence (Supplementary Fig. 15a, b). We also established LS174T-cont cells using a retroviral vector with a target sequence for EGFP, or an empty retroviral vector, and A549-cont cells using an empty retroviral vector.

### Mice

Animal experiment protocols were approved by the Committee for Ethics in Animal Experimentation (approved protocol No. T17-043), and experiments were conducted in accordance with Guidelines for Animal Experiments of the National Cancer Center. C57BL/6J WT mice were obtained from CLEA Japan (Tokyo, Japan). Mieap-knockout (Mieap^−/−^) mice were generated using the Cre/loxP recombination system, as previously reported^32^. Briefly, floxed and trapped alleles were generated using a single construct bearing a gene-trap cassette doubly flanked by LoxP and FRT, located between exons 5 and 8 of the mouse Mieap gene, which is located on chromosome 5. Mieap homozygous (Mieap^−/−^) deficient mice were generated by mating breeding pairs of Mieap heterozygous (Mieap^+/−^) mice. All mice were housed at 22 ± 2°C with a 12 h light/dark cycle with free access to food, CE-2 (CLEA Japan), and water.

### Plasmid construction

#### Constructs containing Mieap

For construction of the plasmid containing N-terminal EGFP-tagged Mieap, the nucleotide sequence of Mieap was PCR-amplified using primers N-EGFP-Mieap-F and N-EGFP-Mieap-R. PCR products were digested with Kpn I and ligated into pEGFP-C1 (Clontech) cut with the same enzyme. For construction of the plasmid containing C-terminal EGFP-tagged Mieap, the nucleotide sequence of Mieap, excluding the stop codon, was PCR-amplified using the primers C-EGFP-Mieap-F and C-EGFP-Mieap-R. PCR products were ligated into the pCR-Blunt II-TOPO vector (Thermo Fisher Scientific) and sequenced. Inserted products were excised using Hind III restriction sites, and ligated into pEGFP-N1 (Clontech), cut with the same enzyme. N-terminal EGFP-tagged Mieap was used as EGFP-Mieap, except for Fig. 1a – d and Supplementary Movie 3 where the C-terminal EGFP-tagged Mieap was used.

Plasmids containing N-FLAG-Mieap (pN-FLAG-Mieap) were constructed as follows. The nucleotide sequence of Mieap was PCR-amplified using the primers, N-FLAG-Mieap-F and N-FLAG-Mieap-R. PCR products were ligated into the pCR-Blunt II-TOPO vector (Thermo Fisher Scientific) and sequenced. Inserted products were excised using the Kpn I restriction sites and ligated into pre-digested pcDNA3.1 (+) (Thermo Fisher Scientific) cut with the same enzyme. The nucleotide sequence of Mieap was excised from the plasmid using the Hind III and Xho I restriction sites, and ligated into pre-digested pCMV-Tag2A (Agilent) cut with the same enzyme.

The plasmid containing C-FLAG-Mieap (pC-FLAG-Mieap) was constructed as follows. The nucleotide sequence of Mieap, excluding the stop codon, was PCR-amplified using the primers, C-EGFP-Mieap-F and C-EGFP-Mieap-R, the same primers used for construction of the plasmid containing C-terminal EGFP-tagged Mieap. PCR products were ligated into the pCR-Blunt II-TOPO vector (Thermo Fisher Scientific) and sequenced. Inserted products were excised using the Hind III restriction site and ligated into pre-digested p3xFLAG-CMV-14 (Sigma Aldrich) cut with the same enzyme.

Prior to construction of plasmids containing EGFP-Mieap deletion mutants, point mutations in Bgl II, Sac I, EcoR I, and Pst I restriction sites of the multiple cloning site of pEGFP-Mieap were introduced using QuikChange Site-Directed Mutagenesis Kits (Agilent) with primers Mut-F1, Mut-R1, Mut-F2 and Mut-R2, which were confirmed by DNA sequencing.

For construction of plasmids containing EGFP-MieapΔCC (pEGFP-MieapΔCC), the nucleotide sequence of pEGFP-Mieap between two Pst I restriction sites was deleted by digestion with Pst I. The remainder was self-ligated, additionally deleting c.810C using the QuikChange Site-Directed Mutagenesis Kit (Agilent) with primers Mut-F3 and Mut-R3 to make the deletion mutation in-frame.

For construction of plasmids containing EGFP-MieapΔ275 (pEGFP-MieapΔ275), the nucleotide sequence of pEGFP-Mieap between the Bgl II and Sma I restriction sites was deleted by digestion using Bgl II and Sma I. After blunting with T4 DNA polymerase (Thermo Fisher Scientific), the remainder was self-ligated.

For construction of plasmids containing EGFP-MieapΔ496 (pEGFP-MieapΔ496), the nucleotide sequence of pEGFP-Mieap was deleted between the EcoR I and Kpn I restriction sites by digestion using EcoR I and Kpn I. After blunting with T4 DNA polymerase (Thermo Fisher Scientific), the remainder was self-ligated.

For construction of plasmids containing TagRFP-T-Mieap (pTagRFP-T-Mieap), the nucleotide sequence of pEGFP-Mieap between the Nhe I and Xho I restriction sites containing EGFP was replaced with nucleotide sequence of pTagRFP-T-EEA1 (Addgene #42635) between the Nhe I and Xho I restriction sites containing TagRFP-T, by digestion using Nhe I and Xho I.

For construction of plasmids containing GST-Mieap (pGST-Mieap), the nucleotide sequence of Mieap (amino acids 99-298) was PCR-amplified using the primers, GST-Mieap-F and GST-Mieap-R. PCR products were ligated into the pCR-Blunt II-TOPO vector (Thermo Fisher Scientific) and sequenced. Products were digested with EcoR I and Xho I, and ligated into pGEX5X-2 (Cytiva).

For construction of plasmids containing Mieap ΔCC (pMieap ΔCC), pEGFP-Mieap ΔCC was digested at the Kpn I restriction sites to obtain the nucleotide sequence of Mieap ΔCC, and ligated into pcDNA3.1 (+) (Thermo Fisher Scientific) cut with the same enzyme, Kpn I.

For construction of plasmids containing Mieap Δ274 (pMieap Δ274), the nucleotide sequence of Mieap Δ274 was PCR-amplified from pEGFP-Mieap Δ275 using the primers, Δ274-F and Δ274-R. PCR products were digested with Kpn I, and ligated into pcDNA3.1 (+) (Thermo Fisher Scientific) cut with the same enzyme, Kpn I.

For construction of plasmids containing Mieap Δ496 (pMieap Δ496), pEGFP-Mieap Δ496 was subjected to inverse PCR using the primers, ΔEGFP-F and ΔEGFP-R to delete the nucleotide sequence of EGFP from pEGFP-Mieap Δ496, and the product was self-ligated using KOD-Plus-Mutagenesis Kit (TOYOBO).

Prior to construction of plasmids containing TagRFP-T-Mieap deletion mutants, the nucleotide sequence of TagRFP-T was PCR-amplified using the primers, TagRFP-T-F and TagRFP-T-R. PCR products were digested with Hind III and EcoR V, and ligated into pcDNA3.1 (+) (Thermo Fisher Scientific) cut with the same enzymes (pcDNA-N-TagRFP). The nucleotide sequence of Mieap was PCR-amplified using the primers, G35-F and G35-R. PCR products were digested with EcoRV and PspOMI, and ligated into pcDNA-N-TagRFP cut with the same enzymes (pG35).

For construction of plasmids containing TagRFP-T-Mieap ΔCC (pTagRFP-T-Mieap ΔCC), pG35 was subjected to inverse PCR using the primers, ΔCC-F and ΔCC-R, and the product was self-ligated using KOD-Plus-Mutagenesis Kit (TOYOBO).

For construction of plasmids containing TagRFP-T-Mieap Δ275 (pTagRFP-T-Mieap ΔCC), pTagRFP-T-Mieap was subjected to inverse PCR using the primers, Δ275-F and Δ275-R, and the product was self-ligated using KOD-Plus-Mutagenesis Kit (TOYOBO).

For construction of plasmids containing TagRFP-T-Mieap Δ496 (pTagRFP-T-Mieap Δ496), pG35 was subjected to inverse PCR using the primers, Δ496-F and Δ496-R, and the product was self-ligated using KOD-Plus-Mutagenesis Kit (TOYOBO).

All primers are listed in Table S1.

#### Other constructs

For construction of plasmids containing EGFP-BNIP3 (pEGFP-BNIP3), plasmids containing FLAG-BNIP3 (pCMV-Tag2B-BNIP3) were constructed in advance. For construction of the pCMV-Tag2B-BNIP3, the nucleotide sequence of BNIP3 was PCR-amplified using the primers, BNIP3-F and BNIP3-R. PCR products were ligated into the pCR-Blunt II-TOPO vector (Thermo Fisher Scientific) and sequenced. Inserted products were digested with EcoR I and Xho I, and ligated into the pre-digested pCMV-Tag2B (Agilent) cut with the same enzyme. The nucleotide sequence of pCMV-Tag2B-BNIP3 was digested at the EcoR I and Xho I restriction sites, and subsequently blunted with T4 DNA polymerase (Thermo Fisher Scientific). pEGFP-C1 (Clontech) was digested with Bgl II, blunted with T4 DNA polymerase, self-ligated, digested with EcoR I and Sma I, and ligated with the fragment of pCMV-Tag2B-BNIP3.

For construction of plasmids containing EGFP-NIX (pEGFP-NIX), plasmids containing FLAG-NIX (pCMV-Tag2B-NIX) were constructed in advance. For construction of the pCMV-Tag2B-NIX, the nucleotide sequence of NIX was PCR-amplified using the primers, NIX-F and NIX-R. PCR products were ligated into the pCR-Blunt II-TOPO vector (Thermo Fisher Scientific) and sequenced. Inserted products were digested with EcoR I and Xho I, and ligated into pre-digested pCMV-Tag2B (Agilent) cut with the same enzyme. The nucleotide sequence of pCMV-Tag2B-NIX was digested at the EcoR I and Xho I restriction sites, and subsequently blunted with T4 DNA polymerase (Thermo Fisher Scientific). pEGFP-C1 (Clontech) was digested with Bgl II, blunted with T4 DNA polymerase, self-ligated, digested with EcoR I and Sma I, and ligated with the fragment of pCMV-Tag2B-NIX.

For construction of the plasmid backbone containing EGFP (pN-EGFP), the nucleotide sequence of EGFP, excluding the stop codon, was PCR-amplified using the primers, N-EGFP-F and N-EGFP-R. PCR products were digested with Hind III and BamH I, and ligated into pcDNA3.1 (+) (Thermo Fisher Scientific) cut with the same enzymes.

For construction of the plasmid backbone containing EGFP (pC-EGFP), the nucleotide sequence of EGFP was PCR-amplified using the primers, C-EGFP-F and C-EGFP-R. PCR products were digested with BamH I and Not I, and ligated into pcDNA3.1 (+) (Thermo Fisher Scientific) cut with the same enzymes.

For construction of plasmids containing EGFP-TAMM41 (pEGFP-TAMM41), the nucleotide sequence of TAMM41, excluding the stop codon, was PCR-amplified using the primers, TAMM41-F and TAMM41-R. PCR products were digested with Nhe I and BamH I, and ligated into pC-EGFP cut with the same enzymes.

For construction of plasmids containing EGFP-PGS1 (pEGFP-PGS1), the nucleotide sequence of PGS1, excluding stop codon, was PCR-amplified using the primers, PGS1-F and PGS1-R. PCR products were digested with Hind III and BamH I and ligated into pC-EGFP cut with the same enzymes.

For construction of plasmids containing EGFP-PTPMT1 (pEGFP-PTPMT1), the nucleotide sequence of PTPMT1, excluding the stop codon, was PCR-amplified using the primers, PTPMT1-F and PTPMT1-R. PCR products were digested with Nhe I and Hind III and ligated into pC-EGFP cut with the same enzymes.

For construction of plasmids containing EGFP-CRLS1 (pEGFP-CRLS1), the nucleotide sequence of CRLS1 was PCR-amplified using the primers, CRLS1-F and CRLS1-R. PCR products were digested with BamH I and Not I and ligated into pN-EGFP cut with the same enzymes.

For construction of plasmids containing EGFP-PLA2G6 (pEGFP-PLA2G6), the nucleotide sequence of PLA2G6, excluding the stop codon, was PCR-amplified using the primers, PLA2G6-F and PLA2G6-R. PCR products were digested with Nhe I and Kpn I and ligated into pC-EGFP cut with the same enzymes.

For construction of plasmids containing EGFP-TAZ (pEGFP-TAZ), the nucleotide sequence of TAZ, excluding the stop codon, was PCR-amplified using the primers, TAZ-F and TAZ-R. PCR products were digested with Nhe I and Kpn I and ligated into pC-EGFP cut with the same enzymes.

For construction of plasmids containing EGFP-PRELI (pEGFP-PRELI), the nucleotide sequence of PRELI, excluding the stop codon, was PCR-amplified using the primers, PRELI-F and PRELI-R. PCR products were digested with Nhe I and BamH I and ligated into pC-EGFP cut with the same enzymes.

For construction of plasmids containing EGFP-LONP1 (pEGFP-LONP1), the nucleotide sequence of LONP1, excluding the stop codon, was PCR-amplified using the primers, LONP1-F and LONP1-R. PCR products were digested with Hind III and BamH I and ligated into pC-EGFP cut with the same enzymes.

For construction of plasmids containing EGFP-PLD6 (pEGFP-PLD6), the nucleotide sequence of PLD6, excluding the stop codon, was PCR-amplified using the primers, PLD6-F and PLD6-R. PCR products were digested with Nhe I and Kpn I and ligated into pC-EGFP cut with the same enzymes.

All primers are listed in Table S1.

### Transfection

For transfection, cells were seeded (2×10^5^ cells/dish) in 35-mm glass bottom dishes. Plasmids (2 μg /dish) were transfected using FuGENE6 transfection reagent (Promega), according to the manufacturer’s instructions.

### Recombinant adenovirus construction

Ad-Mieap was derived from viruses as previously reported^29, 35, 72^. Replication-deficient recombinant viruses Ad-EGFP-Mieap, Ad-N-FLAG-Mieap, Ad-C-FLAG-Mieap, Ad-EGFP-MieapΔCC (Δ104-270), Ad-EGFP-MieapΔ275, Ad-EGFP-MieapΔ496, Ad-TagRFP-T-Mieap, Ad-mApple-TOMM20 derived from mApple-TOMM20-N-10 (Addgene #54955), Ad-EGFP-BNIP3, Ad-EGFP-NIX, Ad-AcGFP1-Mito, and Ad-DsRed2-Mito were generated from the corresponding plasmid vectors and purified as described previously^73^. Briefly, DNA fragments obtained by restriction of each plasmid vector were blunted using T4 DNA polymerase, ligated into the SmiI site of the cosmid, pAxCAwtit (Takara), which contains the CAG promoter and the entire genome of type 5 adenovirus, except the E1 and E3 regions. Recombinant adenoviruses were generated by *in vitro* homologous recombination in the 293 cell line with the cDNA-inserted pAxCAwtit and the adenovirus DNA terminal–protein complex. Viruses were propagated in the 293 cell line and purified by two rounds of CsCl density centrifugation. Viral titers were determined with a limiting dilution bioassay using 293 cells.

Infection of cell lines was carried out by adding viral solutions to cell monolayers, incubating them at 37°C for 120 min with brief agitation every 20 min. This was followed by addition of culture medium and return of the infected cells to the 37°C incubator.

### Immunocytochemistry

For immunocytochemistry, cells were grown on 8-well chamber slides (1-4×10^4^ cells/well) at 37°C in conventional culture medium, and fixed in paraformaldehyde (Supplementary Fig. 1b, 2%; Supplementary Fig. 1c, 4%) for 15 min at room temperature. Slides were incubated with Triton X-100 (Supplementary Fig. 1b, 0.1% for 2 min; Supplementary Fig. 1c, 0.5% for 10 min), and washed 3x with phosphate-buffered saline (PBS) at room temperature. Cells were blocked with 3% bovine serum albumin (BSA) in PBS (Supplementary Fig. 1b, for 3 h; Supplementary Fig. 1c, for 2 h), and sequentially incubated with rabbit polyclonal anti-Mieap antibody (1:200), mouse monoclonal anti-GFP antibody (1:200), or mouse monoclonal anti-FLAG antibody (1:1000) for 2 h at room temperature. After washing 3x with PBS, slides were incubated with Alexa Fluor 546 goat anti-rabbit IgG antibody (1:200) or Alexa Fluor 546 goat anti-mouse IgG antibody (1:200) at room temperature (Supplementary Fig. 1b, for 2 h; SSupplementary Fig. 1c, for 1 h). Slides were washed 3x with PBS. Then they were mounted with VECTASHIELD H-1000 (Vector Laboratories) and observed using a FLUOVIEW FV3000 confocal laser scanning microscope (Olympus).

### Transmission electron microscopy (TEM)

A549-cont and U373MG cells (4×10^4^ cells/24-well plate) were infected with Ad-Mieap. On day 1 after infection, cells were fixed in phosphate buffered 2.5% glutaraldehyde and subsequently post-fixed in 1% OsO_4_ at 4°C for 2 h. Then, specimens were dehydrated in a graded ethanol series and embedded in epoxy resin. Ultrathin sections (75 nm) were cut with an ultramicrotome. Ultrathin sections stained with uranyl acetate and lead staining solution were observed on a transmission electron microscope H-7500 (Hitachi) at 80 kV.

LS174T (control and Mieap-KD) cells cultured under normal conditions were also processed for TEM as mentioned above, with the following modifications. 2% glutaraldehyde was used for prefixation. 2% OsO_4_ was used instead for post-fixation. 80-90-nm ultrathin sections were cut and observed on a transmission electron microscope H-7600 (Hitachi) at 100 kV.

Kidney and liver specimens were collected from an 18-week-old WT mouse and a 16-week-old Mieap^−/−^ male mouse. BAT specimens were collected from a 40-week-old WT and a 40-week-old Mieap^−/−^ male mouse. Specimens cut into approximately 3 × 3 × 3 mm^3^ were also processed for TEM and processed in the same fashion as the aforementioned A549-cont cells. However, a mixture of 2% paraformaldehyde and 2% glutaraldehyde was used for prefixation.

### Post-embedding immunoelectron microscopy

A549-cont cells (2×10^5^ cells/35-mm glass bottom dish) were infected with Ad-Mieap. On day 1 after infection, cells were fixed with 4% paraformaldehyde and 0.025% glutaraldehyde in 0.1 M PBS (pH 7.4) for 1 h at 4°C. After fixation, cells were washed with 0.1 M PBS (pH 7.4) for 16 h at 4°C, dehydrated in a graded ethanol series, and infiltrated with LR White resin. Polymerization was performed in TAAB embedding capsules (TAAB) inverted on glass-bottom dishes for 3h at 60°C. Ultrathin sections (75 nm) were collected on nickel grids. After blocking with 3% BSA in PBS for 1h, sections were incubated with anti-Mieap antibody (1:200) diluted in PBS with 0.05% Triton X-100 for 2h at RT. Sections were washed 8x with 0.15% glycine in PBS, and incubated with goat anti-rabbit IgG 10-nm gold antibody (1:50) diluted in PBS with 0.05% Triton X-100 for 2h at RT. Sections were washed 8x in PBS and fixed with 1% glutaraldehyde in PBS for 5 min. Sections were washed 8x in distilled water. Grids were embedded in a mixture containing 2.7% polyvinyl alcohol and 0.3% uranyl acetate. Sections on grids were observed on a transmission electron microscope H-7500 (Hitachi) at 75 kV.

### Amino acid sequence analyses of Mieap protein

We analyzed the phylogenetic spread of Mieap orthologs using OrthoDB v10 (https://www.orthodb.org/)37. Multiple sequence alignment for Mieap orthologs was performed using Genetyx ver. 10. Prediction of IDRs in the amino acid sequence of Mieap was done using VL3-BA^39^ on the PONDR server (http://www.pondr.com/) and collated with meta-prediction of IDRs using DisMeta (http://www-nmr.cabm.rutgers.edu/bioinformatics/disorder/)38. Prediction of coiled-coil regions was done using COILS (https://embnet.vital-it.ch/software/COILS_form.html)40. Hydrophobicity of Mieap was analyzed according to the Kyte-Doolittle index^42^ using ProtScale (https://web.expasy.org/protscale/)74. The linear net charge per residue of Mieap was analyzed using CIDER (http://pappulab.wustl.edu/CIDER/)41.

### Analyses of confocal microscopy image data

Throughout the study, confocal microscopy images were taken with a FLUOVIEW FV3000 confocal laser scanning microscope (Olympus). For validation of the spatial relationship between Mi-BCs and mApple-TOMM20, additional images were taken using a SpinSR10 spinning disk confocal super resolution microscope (Olympus). For Z-stack and time-lapse imaging, a montage of differential interference contrast (DIC) and fluorescence images was created using MetaMorph ver. 7.8 (Molecular Devices). 3D reconstruction was performed using cellSens Imaging Software (Olympus). Line-scan profiles were acquired using MetaMorph ver. 7.8 (Molecular Devices).

### FRAP experiments

EGFP-Mieap, EGFP-MieapΔCC, EGFP-MieapΔ275, and EGFP-MieapΔ496 were expressed in A549-cont cells to generate condensates by infection with Ad-EGFP-Mieap, Ad-EGFP-MieapΔCC, Ad-EGFP-MieapΔ275, and Ad-EGFP-MieapΔ496, respectively. FRAP experiments were performed on a FLUOVIEW FV3000 confocal laser scanning microscope (Olympus), using a 60x/1.4 NA oil immersion objective (Olympus). Condensates were subjected to spot-bleaching or full-bleaching (bleaching entire condensates). For spot-bleaching, the bleaching area was unified to a diameter of 1.38 µm. Condensates were imaged for 6 s, acquiring 30 images prior to spot-bleaching or 50 s, acquiring 5 images prior to full-bleaching. Photobleaching employed a 488-nm laser at 10% laser power with 11.6 µs/µm exposure time or 1.4% laser power with 1.4 µs/µm exposure time. Time-lapse images were acquired at 0.2-ms intervals for 60 s or 10 s intervals for 15 min. Spot-bleaching data for each construct were acquired from 15 different condensates. Full-bleaching data of each construct were acquired from 10 different condensates.

### Calculation of intensity ratio

To evaluate partitioning of EGFP-Mieap and deletion mutant proteins, EGFP-Mieap WT, ΔCC, Δ275, and Δ496 were expressed in A549-cont cells to generate condensates by infection with Ad-EGFP-Mieap WT, ΔCC, Δ275, and Δ496, respectively. EGFP intensity of condensates and cytoplasm was measured. Because EGFP intensity of these condensates was higher than the intensity of 0.4 mg/mL His-EGFP solution for standard curve, we chose intensity ratio rather than partition coefficient for the parameter of this partitioning experiments^52^. Intensity ratio was calculated as (Intensity of condensates-Intensity of background)/(Intensity of cytoplasm-Intensity of background), where Intensity of condensates, Intensity of cytoplasm, and Intensity of background are the mean intensities of condensates, cytoplasm, and PBS acquired by the identical conditions (laser wavelength, 488 nm; laser transmissivity, 0.01%; detection wavelength, 500–600 nm; voltage, 350 V) on a FLUOVIEW FV3000 confocal laser scanning microscope (Olympus). Intensity ratio data were obtained from 40 cells for each construct.

### Expression and purification of GST and GST-Mieap

*Escherichia coli* (BL21) cells transformed with expression vectors were grown in 200 mL of Luria-Bertani medium at 37°C until the OD600 was between 0.55-0.6. Protein expression was induced with 100 µM IPTG, and bacteria were subsequently incubated for 3 h at 25°C. After harvesting bacteria by centrifugation at 3000 × *g* for 10 min at 4 °C, pellets were lysed with lysis buffer (1% Triton X-100 buffered in PBS supplemented with 1 mM Phenylmethylsulfonyl fluoride), and sonicated (20 × 30 s bursts with 10 s rest between bursts). Insoluble material was removed by centrifugation at 10,000 rpm for 30 min at 4 °C. Supernatant was incubated with glutathione-Sepharose 4B (Cytiva) pre-equilibrated with lysis buffer at 4°C overnight. After the beads were washed twice with lysis buffer, proteins were eluted with elution buffer (50 mM glutathione diluted in 50 mM Tris–HCl, pH 8.0), and dialyzed at 4°C overnight against PBS.

### Lipid-binding analysis

For lipid-binding analysis, protein-lipid interactions on lipid-spotted membranes were evaluated with fat blot assays^49^. Natural CL, PC, and PE derived from bovine heart (Olbracht Serdary Research Laboratories) were diluted with chloroform/methanol/1N HCl (80:80:1). 1 μL of each diluted lipid was spotted onto PVDF membranes (Cytiva) for antigen-antibody reactions using anti-Mieap antibody or nitrocellulose membranes (Cytiva) for antigen-antibody reactions using an anti-GST antibody to align spots with increasing amounts of lipids ranging from 0-667 pmol. Here, approximate molarities of CL, PC, and PE calculated from molecular weights of tetralinoleoyl CL, distearoyl PC, and distearoyl PC were used, respectively. After membranes were blocked with blocking buffer (3% fatty acid-free BSA diluted in 50 mM Tris–HCl, 150 mM NaCl, pH 7.5) for 1 h, membranes were incubated with 2.5 μg/mL of GST-Mieap or GST protein diluted in blocking buffer containing 0.1% Tween 20 overnight. Membranes were incubated with primary antibody (rabbit anti-Mieap antibody or rabbit anti-GST antibody) diluted in blocking buffer containing 0.06% Tween 20 (1:1000) for 3.5 h, and subsequently a secondary antibody (goat anti-rabbit antibody conjugated to horseradish-peroxidase) diluted in blocking buffer containing 0.06% Tween 20 (1:10000) for 1 h. ECL Western Blotting Detection Reagents (Cytiva) was used to detect HRP and chemiluminescence was visualized with an ImageQuant LAS 4000 system (Cytiva).

### Lipid preparation

Lipid preparation was performed as described previously^75, 76^. Briefly, total lipids were extracted from samples using the Bligh-Dyer method^77^. An aliquot of the organic phase was added to an equal volume of methanol before being loaded onto a DEAE-cellulose column (Wako Chemical) pre-equilibrated with chloroform. After successive washes with chloroform/methanol (1:1, v/v), acidic phospholipids were eluted with chloroform/methanol/HCl/water (12:12:1:1, v/v), followed by evaporation to dryness to yield a residue was soluble in methanol.

### Mass spectrometric analyses of CL

Analyses were performed on an LC/MS/MS system consisting of a Q-Exactive Plus mass spectrometer (Thermo Fisher Scientific) equipped with an electrospray ionization source and an UltiMate 3000 system (Thermo Fisher Scientific). Lipid samples were separated on a Waters X-Bridge C_18_ column (3.5 μm, 150 mm × 1.0 mm i.d.) at 40°C using a solvent step-gradient as follows: mobile phase A (isopropanol/methanol/water (5:1:4, v/v/v) supplemented with 5 mM ammonium formate and 0.05% ammonium hydroxide (28% in water))/mobile phase B (isopropanol supplemented with 5 mM ammonium formate and 0.05% ammonium hydroxide (28% in water)) ratios of 60%/40% (0 min), 40%/60% (1 min), 20%/80% (9 min), 5%/95% (11-30 min), 95%/5% (31-35 min) and 60%/40% (45 min). Flow rate was 25 μL/min. Source and ion transfer parameters applied were as follows. Spray voltage was 3.0 kV. For negative ionization modes, the sheath gas and capillary temperatures were maintained at 60 and 320 °C, respectively. The Orbitrap mass analyzer was operated at a resolving power of 70,000 in full-scan mode (scan range: 200–1800 m/z; automatic gain control (AGC) target:3e6) and of 35,000 in the Top 20 data-dependent MS2 mode (stepped normalized collision energy: 20, 30 and 40; isolation window: 4.0 m/z; AGC target: 1e5). Identification of CL molecular species was performed using LipidSearch 4.2 software (Mitsui Knowledge Industry).

### Real-time ATP rate assay

LS174T-cont and Mieap-KD cells were seeded at a density of 2.5×10^4^ cells/well (n=9) on a Seahorse XF24 Cell Culture Microplate. Cells were incubated at 37°C in a humidified chamber with 5% CO_2_. 18 h after seeding, culture medium was replaced with XF DMEM medium pH 7.4 supplemented with 25 mM glucose and 2 mM L-glutamine through three washes.

HCT116 cells were seeded at a density of 0.8×10^6^ cells/60-mm dish (n=9). Cells were incubated at 37°C in a humidified chamber with 5% CO_2_. 24 h after seeding, cells were treated with Ad-Mieap or Ad-empty. 24 h after infection, cells were reseeded at a density of 4×10^4^ cells/well (n=9) on a SeahorseXF24 Cell Culture Microplate. 20 h after reseeding, culture medium was replaced with XF DMEM medium pH 7.4 supplemented with 25 mM glucose and 2 mM L-glutamine through three washes.

After cells were incubated at 37 °C in a non-CO_2_ incubator for 60 min, cell culture plates were loaded into a Seahorse XFe24 Analyzer. Oxygen consumption rate (OCR) and extracellular acidification rate (ECAR) were recorded before and after serial injections of oligomycin and rotenone/antimycin A to yield final concentrations of 0.5 μM.

### Flow cytometric analysis

LS174T-cont and Mieap-KD cells cultured under normal conditions were harvested by trypsin-EDTA treatment. After adding complete growth media to inactivate trypsin, cells were centrifuged, washed with PBS, and incubated with 5 μM 2′,7′-dichlorofluorescin-diacetate (Sigma) for 20 min at 37°C. After being washed with PBS, cells were immediately analyzed with an EC800 flow cytometry analyzer (Sony) using the 488-nm line.

### Primers

The information of all PCR primers is indicated in Table S1.

### Quantification and statistical analysis

#### FRAP data quantification

Fluorescence recovery rates were calculated using cellSens Imaging Software (Olympus), in which the intensity initially acquired after bleaching was set to 0 and the pre-bleaching intensity was set to 1. The normalized average fluorescence recovery was plotted in JMP 14.2.0 (SAS).

#### Crista data quantification

For quantification of crista data, crista area and outlines of mitochondrial sections in TEM images were marked manually using Adobe Photoshop CC, where normal crista morphology was identified by the presence of lamellar structures with distinct OsO_4_ staining. Aberrant crista-like structures that were not observed in mitochondria of WT were excluded. Subsequently, the ratio of crista area per mitochondrial section was calculated from the indicated number of mitochondria in legends of Fig. 8D, L, M, and 9F, using Image J^78^.

#### Statistical analysis

Statistical analyses were performed in JMP 14.2.0 (SAS). Levels of significance in Figure 6I, 8A, 8B, 8D, 8H–J, 8L, 8N, 9B–D, 9F, S10B–D, S10F–H were assessed using Student’s two-tailed t-tests. Levels of significance in mass spectrometric analyses for biological replicate pairs shown in Figure 5A, 5B, 8E, and 8Ff were assessed using the paired two-tailed t-test. p < 0.05 was considered statistically significant. Asterisks were allotted to all the Figures containing statistical analyses as follows: *, p < 0.05; **, p < 0.01, ***, p < 0.001, ****, p < 0.0001.

### Data visualization

Visualization of the experimental data subjected to statistical analyses were performed using Graph Builder engine in JMP 14.2.0 (SAS). When the data were visualized using violin plots, box plots were overlaid. The center line in the box indicates the median. The bottom and top of the box indicate the 25^th^ and 75^th^ percentiles. The whiskers extend 1.5 times the interquartile range (IQR) from the top and bottom of the box unless the minimum and maximum values are within the IQR. The values which fall above or below the whiskers are plotted individually as outliers.

### Data and code availability

The datasets generated during the mass spectrometric analyses of cardiolipin are available in the Metabolomics Workbench repository, [https://www.metabolomicsworkbench.org/data/DRCCMetadata.php?Mode=Project&ProjectID=PR001192], under Project ID PR001192. All other relevant data which support the findings of this study are included in this article and its supplementary information files. The codes used for data analysis are available from the corresponding author upon a request.

## Supplemental information

### ● Supplemental tables

Table S1. Oligonucleotides used in this paper, related to STAR methods

### ● Supplementary figures

Figure S1 (related to Figure 1). Mieap forms biomolecular condensates

(**A**) Comparative imaging of EGFP-Mieap condensates, visualized with both EGFP-Mieap and immunofluorescence (IF) using anti-Mieap antibody (upper panel) or anti-GFP antibody (lower panel) in A549 cells.

(**B**) IF imaging of N-FLAG-Mieap condensates (upper panel) or C-FLAG-Mieap condentates (lower panel) using anti-FLAG antibody in A549 cells.

(**C**) Transmission electron microscopy of Mieap condensates stained with osmium (OsO_4_).

(**D**) Post-embedding immunoelectron microscopy of Mieap condensates using anti-Mieap antibody.

Figure S2 (related to Figure 2). Analyses of Mieap orthologs

Amino acid sequence analyses of representative Mieap orthologs, as in Figure 2E.

Figure S3 (related to Figure 3). FRAP analysis of condensates formed by EGFP-Mieap and three deletion mutants

(**A**) Representative FRAP images of condensates formed by EGFP-Mieap (WT) and three deletion mutants (ΔCC, Δ275, and Δ496) in A549 cells. Each condensate was subjected to spot-bleaching using a 488-nm laser at 10% laser power with an 11.6 μs/μm exposure time and followed up for 60 s. Bleached areas are indicated by yellow arrows. Scale bar, 2 μm.

(**B**) Plotting of normalized average fluorescence recovery in the FRAP experiment with weaker laser exposure. Laser power was weakened to 1.4% and the exposure time was shortened to 1.4 μs/μm. n = 15 condensates for each construct. Data shown are means ±SD.

Figure S4 (related to Figure 4). Verification of subcellular localization of fluorescently labeled proteins

(**A** – **I**) Subcellular localization of EGFP-TAMM41 (**A**), EGFP-PGS1 (**B**), EGFP-PTPMT1 (**C**), EGFP-CRLS1 (**D**), EGFP-PLA2G6 (**E**), EGFP-TAZ (**F**), EGFP-PRELI (**G**), EGFP-LONP1(**H**), and EGFP-PLD6 (**I**) verified by confocal live cell imaging, compared with localization of MitoTracker Red in A549 cells.

Figure S5 (related to Figure 6). Reactive oxygen species (ROS) levels increase in Mieap-KD cells

ROS levels of LS174T-cont and Mieap-KD cells cultured under normal conditions, analyzed by flow cytometry using 2′,7′-dichlorofluorescein-diacetate. Data from duplicate experiments are shown.

Figure S6 (related to Figure 7). Mieap prevents obesity

Body weights of 1,225 Mieap^+/+^, Mieap^+/−^, and Mieap^−/−^ mice (n = 315 Mieap^+/+^, 571 Mieap^+/−^, and 339 Mieap^−/−^ mice, 7-130 weeks of age) were weighed. Dots and quadratic regression curves with 95% confidence intervals are shown for each genotype.

Figure S7 (related to Figure 7). Hypothetical models for Mieap-mediated sequential enzymatic reactions in CL metabolism

(**A**) Hypothetical model of biosurfactant activity of Mieap. Mieap (green) exists in the Mieap-containing phase (lipid phase) (black), as a “scaffold” protein and/or as a potential “biosurfactant.” At the boundary between the surfaces of Mi-BCs (aqueous phase) and the Mieap-containing phase (lipid phase) or between the Mieap-containing phase (lipid phase) and the Mieap-depeleted phase (aqueous phase), the hydrophilic N-terminal end of Mieap always faces the aqueous phase at the boundary. (**B**, **C**) Hypothetical model for Mieap-mediated sequential enzymatic reactions in CL metabolism. Black areas indicate the Mieap-containing phase (lipid phase) containing CL and Mieap. Gray areas indicate the Mieap-depeleted (aqueous phase) containing enzymes. Sequential reactions occur at the interface between the surface of Mi-BCs (aqueous phase) and the Mieap-containing phase (lipid phase) (**B**) or between the Mieap-containing phase (lipid phase) and the Mieap-containing phase (aqueous phase) (**C**). Once Mieap (green) stably interacts with PA via its C-terminal region, one of the enzymes transiently and weakly interacts with the N-terminal region of Mieap. When Mieap interacts with TAMM41, PA is converted to CDP-DG. Such reactions between biosynthetic enzymes and corresponding substrates could be repeated until mature CL is produced. Concentration of enzymes and substrates at Mi-BC surfaces, segregation of enzymes and substrates into distinct sub-compartments of Mi-BCs, interfacial catalysis, and biosurfactant activity of Mieap may enable efficient sequential reactions for CL metabolism. Abbreviations: PA, phosphatidic acid; CDP-DG: cytidine diphosphate diacylglycerol; PGP, phosphatidylglycerophosphate; PG, phosphatidylglycerol; CLOX, oxidized cardiolipin; CLN, nascent cardiolipin; CLM, mature cardiolipin.

### ● Supplementary Movies

Video S1 (related to Figure 1). Time-lapse imaging of the self-fusing Mi-BCs

The condensates comprised of EGFP-Mieap were examined in the A549-cont cell.

Video S2 (related to Figure 1). Z-stack, 3D, and time-lapse imaging show mitochondrial localization of Mi-BCs

The spatial relationship between Mi-BCs and mApple-TOMM20 was examined in the A549-cont cells.

Video S3 (related to Figure 3). Z-stack and time-lapse imaging for screening molecules involved in phase separation by Mi-BCs

EGFP-BNIP3, EGFP-NIX, AcGFP1-Mito, DsRed2-Mito, and SYBR Green I were not phase-separated by Mi-BCs.

Video S4 (related to Figure 3). Z-stack and/or time-lapse imaging for phase-separation of NAO by Mi-BCs

Cardiolipin (CL) visualized with 10-nonylacridine orange bromide (NAO) was phase-separated by Mi-BCs in the A549-cont cells.

Video S5 (related to Figure 4). Z-stack and/or time-lapse imaging for phase-separation of CL synthetic and remodeling enzymes by Mi-BCs

Whether EGFP-tagged TAMM41, PGS1, PTPMT1, CRLS1, PLA2G6, TAZ, PRELI, LONP1, and PLD6 are phase-separated by Mi-BCs were examined in the A549-cont cells.

Video S6 (related to Figure 5). 3D reconstruction showing the spatial relationship between Mi-BCs (TagRFP-T-Mieap), ΔCC-BCs (TagRFP-T-ΔCC), Δ275-BCs (TagRFP-T-Δ275), or Δ496-BCs (TagRFP-T-Δ496) and mitochondrial inner membranes visualized with EGFP-TAMM41 in the HeLa cells

Mi-BCs, ΔCC-BCs, and Δ496-BCs are localized in mitochondria whereas Δ275-BCs are detected in the outside of mitochondria.

Video S7 (related to Figure 5). 3D reconstruction showing the spatial relationship between Mi-BCs (TagRFP-T-Mieap), ΔCC-BCs (TagRFP-T-ΔCC), Δ275-BCs (TagRFP-T-Δ275), or Δ496-BCs (TagRFP-T-Δ496) and mitochondrial inner membranes visualized with NAO in the HeLa cells

Mi-BCs, ΔCC-BCs, and Δ496-BCs are localized in mitochondria whereas Δ275-BCs are detected in the outside of mitochondria.

Video S8 (related to Figure 5). 3D reconstruction showing three patterns of Video S6 and Video S7, including BCs, mitochondria, and merge

Mi-BCs, ΔCC-BCs, and Δ496-BCs are localized in mitochondria whereas Δ275-BCs are detected in the outside of mitochondria.

