## Supplementary figures and images for "Mieap forms membrane-less organelles to compartmentalize and facilitate cardiolipin metabolism"

### Supplementary Figure S1

Supplemental Figure S1

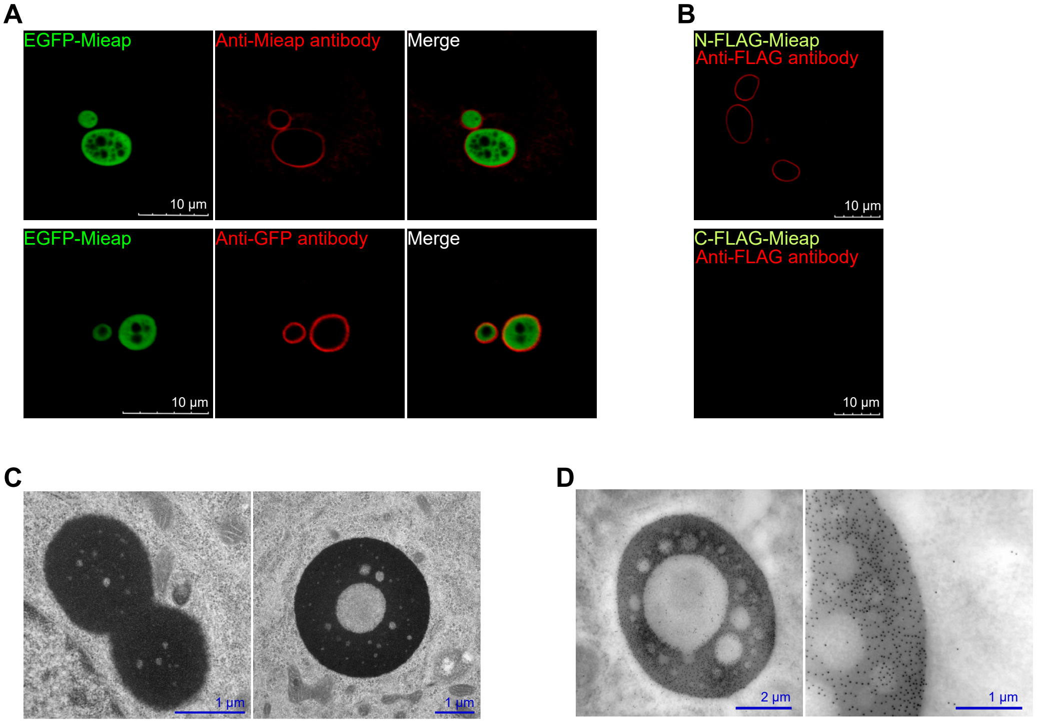

### Supplementary Figure S2

Supplemental Figure S2

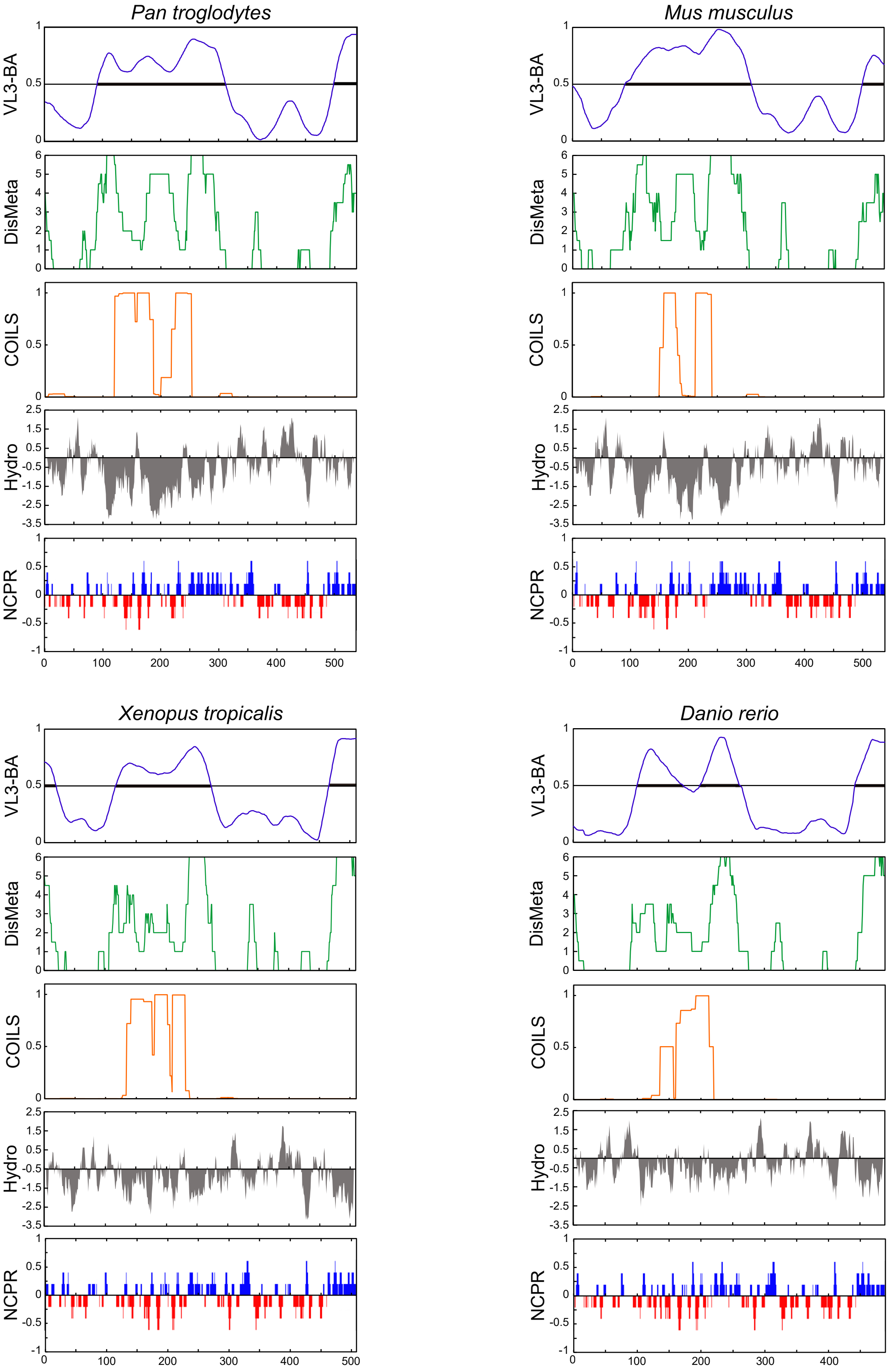

### Supplementary Figure S3

Supplemental Figure S3

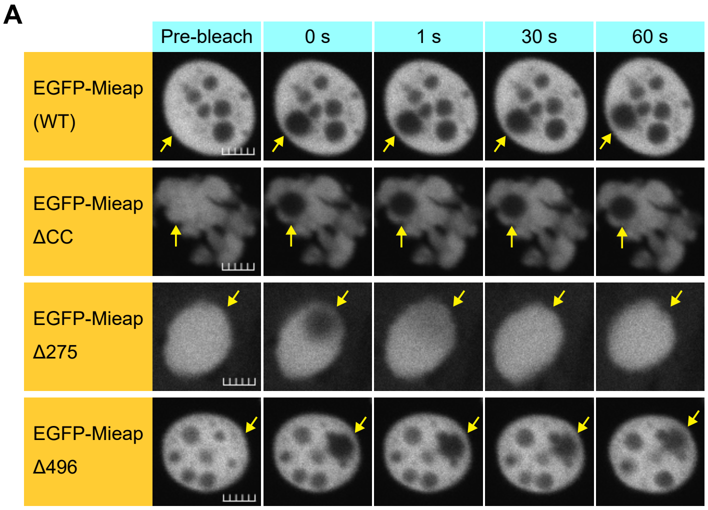


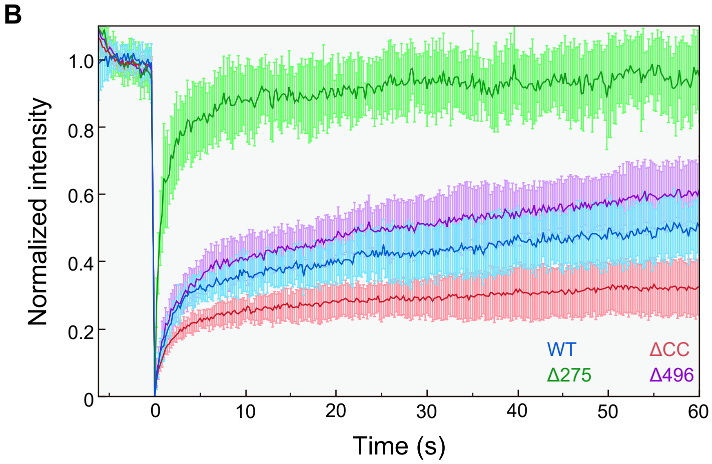

### Supplementary Figure S4

Supplemental Figure S4

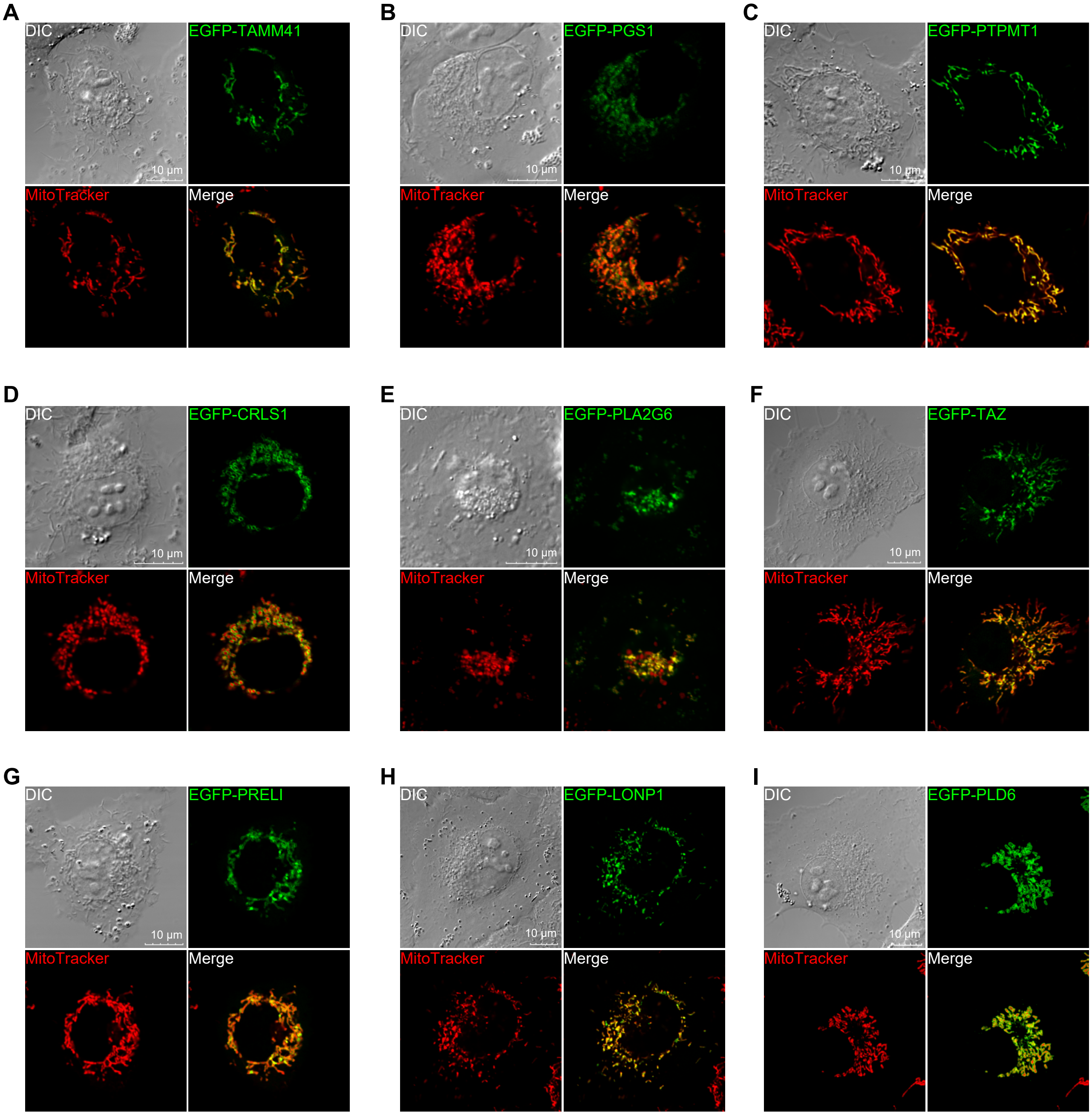

### Supplementary Figure S5

Supplemental Figure S5

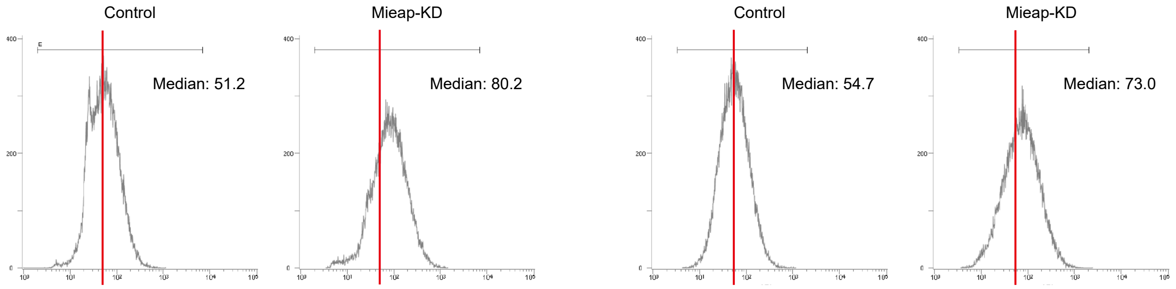

### Supplementary Figure S6

Supplemental Figure S6

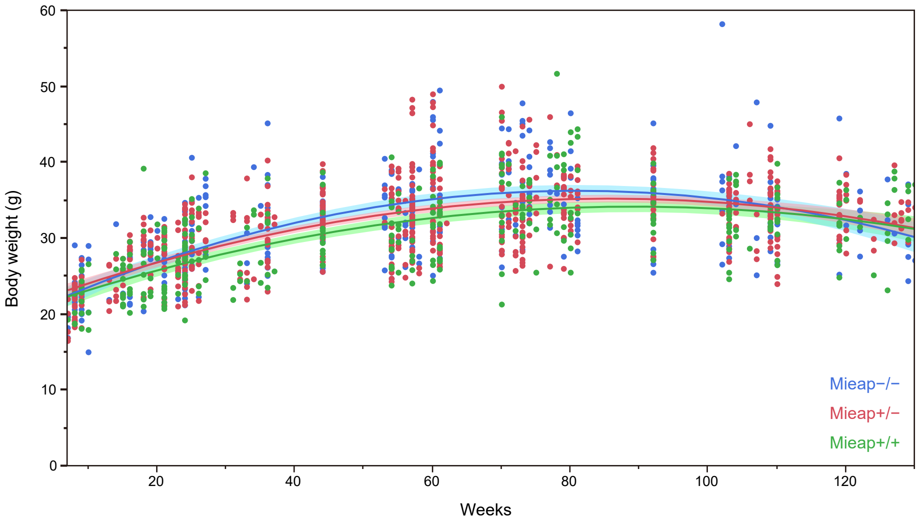

### Supplementary Figure S7

Supplemental Figure S7

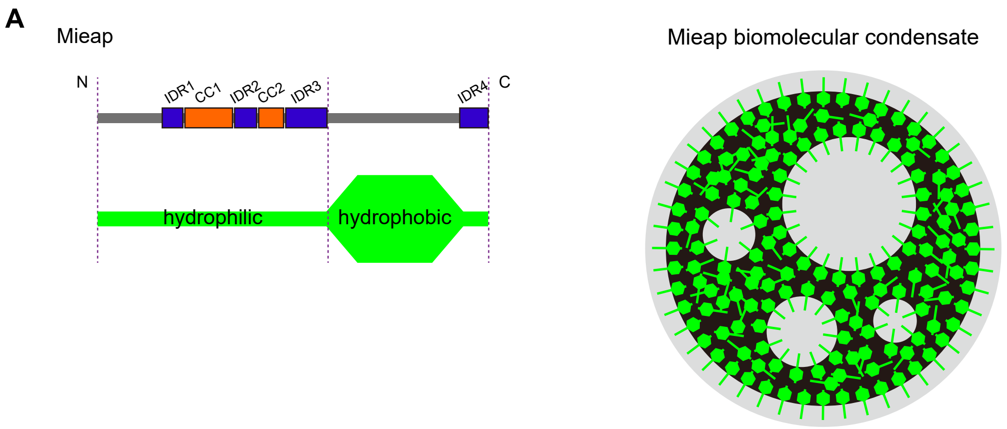


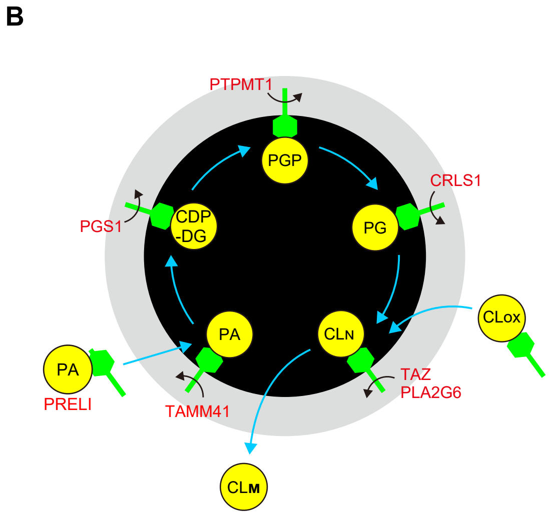


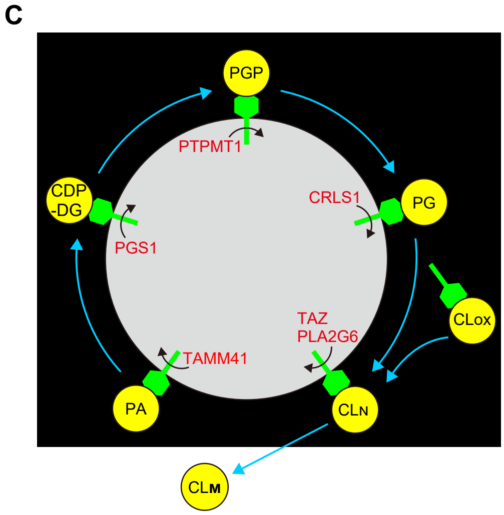
